# MGIDI selection and machine learning reveal harvest index driving traits in sodium azide induced rice mutants with SSR-based genetic diversity

**DOI:** 10.64898/2026.02.17.706299

**Authors:** S. M. Abdullah Al Mamun, Md Rezve, Md. Borhan Ali Sorker, Md. Musabbir Hossain Shoun, Mst. Sabiha Sultana, Aninda Arnab Pandit, Joyanti Ray, Md. Monirul Islam

**Affiliations:** Faculty Member, Agrotechnology Discipline, Khulna University, Khulna-9208, Bangladesh; Post-Graduate Student, Agrotechnology Discipline, Khulna University, Khulna-9208, Bangladesh; Ex-Student, Agrotechnology Discipline, Khulna University, Khulna-9208, Bangladesh

**Keywords:** Sodium azide, M_3_ mutants, SSR markers, Genetic diversity, Multi-trait selection, Machine learning

## Abstract

Sodium azide mutagenesis offers a powerful approach to generate genetic diversity for rice improvement, yet comprehensive characterization of mutant populations using integrated modern breeding tools remains limited. M₃ mutants of BRRI dhan28 induced with sodium azide, were evaluated for 17 agronomic traits and genetic diversity was characterized using 30 SSR markers. The MGIDI was used to characterize elite genotypes and machine learning approaches were used to dissect trait architecture underlying harvest index. The phenotypic variation captured by principal component analysis was 52.12%, and yield was the trait with the highest genotypic variance (278.22) and genotypic coefficient of variation (29.07%). MGIDI analysis detected 10 elite mutants that significantly outperformed within the same environment in combined yield and harvest index. The main predictors of harvest index variability were examined using a Random Forest analysis, and this showed that grain and straw yield were the main predictors of harvest index variability. The SSR markers showed high level of genetic diversity (PIC = 0.264), population structure analysis revealed two subgroups (Fst = 0.0437) and the pairwise genetic distance ranged from 0.000 to 0.733. Procrustean alignment showed a high correlation between molecular and phenotypic variation. An integrated approach of MGIDI selection and prediction of diversity using machine learning underpinned the identification of elite mutants that can be quickly forwarded to breeding programs. This study provides valuable genetic resources and demonstrates that sodium azide mutagenesis combined with modern analytical tools accelerates genetic gains in rice improvement.

## Introduction

Global food security faces unprecedented challenges from the convergence of rapid population growth and climate change. Rice (*Oryza sativa* L.) the second most widely eaten cereal following maize is a major food staple of more than half of the world’s population, especially in Asia, Africa and Latin America (Bandumula, 2018). However, rice production systems are becoming more and more limited by diminishing availability of arable lands contrasting to population pressure, lack of water resources; stagnation in potential yield level and further stressful climatic changes including extremes leading toward low productivity but high unsustainability (Seck et al., 2012). These problems highlight the importance of hastening genetic improvement in rice and breeding climate-resilient high-yielding crops that can maintain productivity under difficult conditions (Ronald, 2014).

In Bangladesh, BRRI dhan28 was officially released in 1994 by the Bangladesh Rice Research Institute (BRRI), and since then has played a major role in ensuring food security of part of its population. Despite being widely cultivated and produced with stable yields for a period spanning more than two decades, BRRI dhan28 now encounters new threats from evolving biological and abiotic stresses as well as continuous demand for yield improvement in response to the increasing food demands. These constraints require constant genetic improvement by modern breeding approaches to keep the variety’s potency and to increase its durability under future rice production systems.

Mutation breeding represents a powerful approach to broaden the genetic base and create novel genetic variation for crop improvement. Unlike traditional hybridization, which is performed based on extant genetic variation within the breeding pool, induced mutagenesis has the potential to produce new alleles and novel phenotypes that are not accessible in natural germplasm (Oladosu et al., 2016). Chemical mutagenesis, especially with sodium azide (SA), has been increasingly adopted as an economic, swift method that highly induces point mutations in plants (Gruszka et al., 2012; Viana et al., 2019). Sodium azide primarily caused G/C to A/T transitions by the production of active organic metabolites and induced a point mutation frequency is about 1.4 - 2.9 Mb across whole genome (Tai et al., 2016). This spectrum of mutational type is especially useful in introducing subtle functional genetic variation which may modulate gene function without inducing a full loss-of-function, and so underpin small effect increments of more complex quantitative traits.

Recent studies have demonstrated that sodium azide can efficiently generate diverse and broad-spectrum blast-resistant mutants, with some mutants displaying durable resistance for nearly two decades (Lo et al., 2022). In addition, sodium azide-induced mutations have been found to revert functional alleles with plant nutritional characteristics like anthocyanin biosynthesis which demonstrate the potential of this mutagen in generating novel genetic variability for breeding programmes (Viana et al., 2019). The M₃ generation, derived from selfing of the M₂ lines is a critical step in mutation breeding programs where segregating heterosis from earlier generations now become significantly fixed offering relative predictability in identifying true-breeding mutants with these desirable traits and allowing for extensive phenotypic and molecular profiling (Oladosu et al., 2016).

Thorough characterization of mutants populations will require concerted use of phenotypic and molecular methods to properly estimate the range and outcomes of induced genetic variation. Phenotypic characterization of agronomic traits is crucial for understanding their expression and potential utilization as breeding materials, but environmental factors among others may cause phenotypic noise that masks genetic differences. Molecular markers, such as Simple Sequence Repeat (SSR) markers provide an alternate tool for the accurate measurement of genetic variation at the DNA level (McCouch et al., 2002). SSR markers are known for their high level of polymorphism, co-dominant nature and average distribution mainly throughout the rice genome, which makes SSR an excellent tool for diversity analysis, population structure estimation and marker-assisted selection (Singh et al., 2024). In rice, SSR markers have been used in recent investigations for estimating genetic diversity of germplasm resources and have provided insights into population structure, phylogenetic relationships and marker-trait associations as useful information for breeding (Hoque et al., 2015).

By integrating multivariate statistical methods with molecular-based diversity analysis, a broader picture of trait architecture and genetic relationships in mutant populations can be obtained. Principal component analysis (PCA), cluster analysis and correlation studies can facilitate understanding of the genetic architecture of complex quantitative traits and in identifying genetically diverse superior lines which could be taken into breeding programs for further improvement (Singh et al., 2015). The recent development of multi-trait selection indices, e.g., the Multi-Trait Genotype-Ideotype Distance Index (MGIDI), also provides a powerful tool for improvement of correlated traits simultaneously by accounting for their correlation structure and intuitive interpretation and allowing simultaneous comparison to an idealized profile of genotype performance (Olivoto and Nardino, 2021; Al Mamun et al., 204). Integrated with machine learning algorithms of Random Forest and Partial Least Squares regression, these approaches can identify the key drivers for complex traits such as Harvest Index and Yield Potential, thereby facilitating genetic gain in breeding programs (Gill et al., 2022).

Although mutation breeding has been successfully conducted for the improvement of rice, characterization of sodium azide-induced mutants is not well explored especially in the M₃ generation of elite varieties such as BRRI dhan28. The lack of this knowledge is impeding efficient applications of chemically induced mutant populations in rice breeding. A comprehensive understanding of diversity, both in morphology and genetics, produced by SA mutagenesis and how this diversity translates into better agronomic performance is important for germplasm selection as well as breeding progress. Therefore, this study was conducted to elucidate the genetic variability, trait architecture and breeding value in 100 M₃ mutant lines of BRRI dhan28 that were treated with sodium azide.

The specific objectives were to: (1) evaluate phenotypic variability across 17 key agronomic traits and assess heritability and genetic advance for selection efficiency; (2) analyze genetic diversity and population structure using 30 polymorphic SSR markers distributed across the rice genome; (3) identify elite mutants with superior harvest index and yield performance using the Multi-Trait Genotype-Ideotype Distance Index (MGIDI); (4) elucidate the trait architecture underlying harvest index variation through machine learning approaches including Random Forest and Partial Least Squares regression; (5) assess phenotypic correlations among yield-contributing traits to understand biological trade-offs and synergies; and (6) integrate phenotypic and molecular data to identify genetically distinct, high-performing mutants suitable for direct utilization in breeding programs or as sources of favorable alleles for marker-assisted introgression. This integrative approach combining classical quantitative genetics, molecular diversity analysis, multi-trait selection indices, and machine learning provides a comprehensive framework for efficient exploitation of mutation-induced genetic variation in rice improvement.

## Materials and methods

### Location and climate of the experimental site

The study was conducted at the Batiaghata, Khulna, Bangladesh (S1 Fig). The soil was characterized as clay loam, exhibiting a pH of 8.1. The experiment was conducted during the Kharif II season, during July to November 2023. Climatic conditions throughout the experimental period were recorded and are presented in S2 Fig. (Data source: Regional Weather Station in Khulna.)

### Plant material, seedling development, and experimental design

The experiment involved 100 M_3_ mutant lines of rice developed through sodium azide mutagenesis, along with the parent variety BRRI dhan28. Seeds were treated with Vitavax-200 (2.5 g kg^-1^) before sowing to prevent seed-borne diseases. Germination was carried out under controlled conditions, and healthy seedlings were transplanted into pots for initial screening. 30 days after sowing, uniform seedlings were transplanted into the field following a Randomized Complete Block Design (RCBD) with three replications. Each mutant line and the parent were grown in individual plots. Standard agronomic practices were followed uniformly to ensure optimal growth and reliable comparison.

### Data collection parameters

Data were recorded for 17 quantitative agronomic traits to evaluate the performance of the M_3_ mutant lines and the parent variety. The parameters included: days to flowering, days to maturity, plant height (cm), panicle length (cm), total tillers hill^-1^, effective tillers hill^-1^, primary and secondary branches panicle^-1^, flag leaf length (cm), grains (filled and total) panicle^-1^, grain length and breadth (mm), 1000-grain weight (g), grain yield hill^-1^ (g), straw yield hill^-1^ (g), and harvest index.

### Multi-trait genotype–ideotype distance index (MGIDI) analysis

Multi-trait selection was performed using the genotype–ideotype distance index (MGIDI) to identify elite mutant genotypes based on the simultaneous improvement of multiple agronomic traits. Prior to analysis, traits were standardized as:

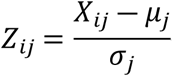 where *X*_*ij*_is the value of genotype *i*for trait *j*, and *μ*_*j*_and *σ*_*j*_are the mean and standard deviation of trait *j*. Trait directions were defined as either higher- or lower-is-better depending on breeding objectives.

Factor analysis was applied to the standardized dataset, and MGIDI for genotype *i*was calculated as the Euclidean distance between the genotype and the ideotype in factor space:

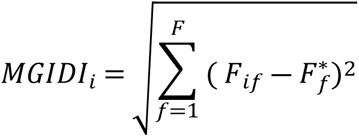 where *F*_*if*_is the score of genotype *i*on factor *f*, and 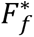represents the ideotype score. Genotypes with the lowest MGIDI values were selected as elite mutants for downstream analyses.

### Random Forest–Based Machine Learning Analysis and Validation

Random Forest (RF) regression was employed to identify key agronomic traits governing harvest index (HI) in rice. Standardized phenotypic traits were used as predictor variables and HI as the response variable. The RF model was implemented using 300 trees with a maximum depth of 5 to capture nonlinear relationships while minimizing overfitting.

The RF prediction was calculated as the mean output of all trees in the ensemble:

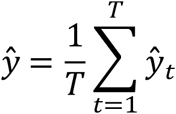

where *ŷ* is the predicted harvest index, *T*is the number of trees, and *ŷ_t_* is the prediction from the *t*-th tree.

Model performance was evaluated using five-fold cross-validation, and predictive accuracy was quantified using the coefficient of determination (R²) between observed and cross-validated predicted HI values. Internal model stability and generalization ability were further assessed using out-of-bag (OOB) estimation inherent to the RF algorithm.

Trait importance was quantified using the mean decrease in impurity (MDI):

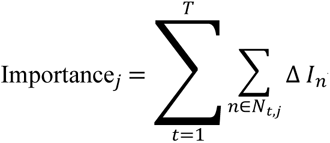 where Δ*I*_*n*_denotes the impurity reduction at node *n*caused by splits on trait *j*, and *N*_*t*,*j*_represents the set of nodes in tree *t*where trait *j*was used. Traits with higher importance values were considered major contributors to HI variation.

### Trait-wise variability and association analysis of yield-contributing traits in MGIDI-selected elite mutants

Phenotypic variability of key yield-contributing traits among the top 10 MGIDI-selected elite rice mutants was evaluated using combined violin–box plots, which depict replication-level distributions, medians, interquartile ranges, and data density to assess trait stability and divergence within elite genotypes. Associations between grain yield per hill (GYH) and individual yield-contributing traits were quantified using Pearson’s correlation and linear regression. Pearson’s correlation coefficient was calculated as:

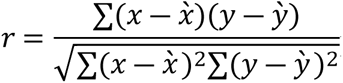 Linear regression was fitted as:

*GHY* = *β*_0_ + *β*_1_*X* + *ε* where *X*represents the explanatory trait. Regression lines with 95% confidence intervals were used to visualize the strength and direction of trait–yield relationships.

### Genomic DNA extraction and SSR marker selection

Young, healthy leaf tissues were collected from each genotype at the age of 21 days for genomic DNA extraction. DNA was isolated using the PureLink Plant Genomic DNA Extraction and Purification Kit (Invitrogen, USA) following the manufacturer’s protocol. The concentration and purity of the extracted DNA were assessed using a Nanodrop 2000 spectrophotometer and their final concentration was adjusted to 30 ng μl^-1^ for PCR reaction. A set of 30 SSR markers was selected from the Gramene database based on their high polymorphism information content (PIC), consistent amplification, and reliable co-dominant allele segregation. The markers used to assess genetic diversity among the M_3_ mutant lines are detailed in S1 Table.

### PCR amplification and gel electrophoresis

PCR amplification was performed in 20 μL reaction volumes containing 50 ng of genomic DNA, 0.5 μM of each primer (forward and reverse), 200 μM of each dNTP, 1.5 mM MgCl_2_, 1× PCR buffer, and 1 unit of Taq DNA polymerase. The amplification followed the protocol of Williams et al. (1990). The thermal cycling conditions included an initial denaturation at 94 °C for 5 minutes, followed by 40 cycles of denaturation at 94 °C for 30 seconds, annealing at marker-specific temperatures (55 °C to 67 °C) for 45 seconds, and extension at 72 °C for 1 minute. A final extension step was carried out at 72 °C for 10 minutes, as described by Chen et al. (1997). Amplified PCR products were separated on 1% agarose gels stained with ethidium bromide. The bands were visualized under UV transillumination and documented following the standard protocol outlined by Sambrook et al. (1989).

### Genetic Diversity Analysis

Genetic diversity Parameters such as number of observed alleles (Na), effective number of alleles (Ne), observed heterozygosity (H_o_) Expected Heterozygosity using Shannon’s information index and fixation index, were calculated in GenAlEx version 6.501 (Peakall and Smouse, 2012). A Nei’s genetic distance (Nei, 1972) among populations was estimated by PowerMarker version 3.25 (Liu and Muse, 2005), and phylogenic relationships were constructed according to the agglomerative hierarchical clustering. Population structure and ancestry of the germplasm collection were analyzed by STRUCTURE software version 2.3.4 (Pritchard et al., 2000). We used a model of admixture with correlated allele frequencies and estimated the best genetic clustering (K), running simulations from K = 2 to K = 9, each over 100,000 iterations after burn-in of 50,000 iterations. The optimal K value was determined using the ΔK method (Evanno et al., 2005).

### Genetic diversity, phenotypic–genotypic concordance, and network-based elite germplasm identification

SSR-based molecular diversity was summarized using Principal Coordinate Analysis (PCoA) based on Jaccard genetic distances, while phenotypic variation was summarized using PCA. Concordance between phenotypic and molecular structures was evaluated using Procrustes analysis:

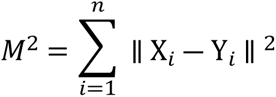Significance was assessed using 999 permutations.

Phenotypic and molecular distance matrices were integrated using equal weights to derive a composite distance:

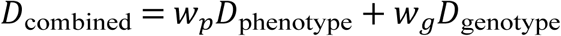

which was converted to a similarity matrix:

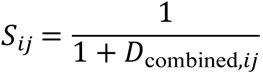 A genotype similarity network was constructed using a high-quantile similarity threshold. Network topology was quantified using degree centrality:

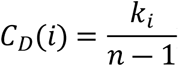 High-yielding genotypes were identified using an upper yield quantile, and those with high centrality were classified as central elite mutants, while low-connectivity but high-yielding genotypes were classified as peripheral elite mutants.

### Statistical analysis

Analysis of variance (ANOVA) was conducted using R software to assess variation among treatments. Means were compared using Duncan’s Multiple Range Test (DMRT). Variance components were estimated to study the genetic and environmental influences on the quantitative traits using the following formulae-

Genotypic variance: 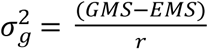 (Burton and Devane, 1953)

Phenotypic variance: 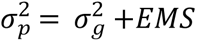 (Burton and Devane, 1953)

Genotypic coefficient of variation: 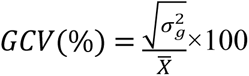 (Johnson et al.,1955)

Phenotypic coefficient of variation: 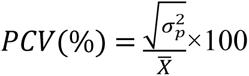 (Johnson et al.,1955)

Broad-sense heritability: 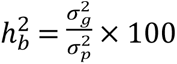 (Johnson et al.,1955)

Genetic advance: 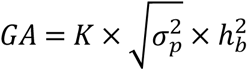 (Johnson et al.,1955)

Genetic advance in percentage of mean: 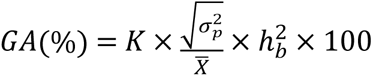 (Comstock and Robinson, 1952).

Where, GMS means the genotypic mean square, EMS means the error mean square, r indicates the number of replications, 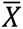 indicates the mean value of the trait, 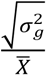 signifies the genotypic standard deviation, 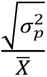 signifies the phenotypic standard deviation, and K is selection differential (2.06 at 5% selection intensity).

Multi-trait selection (MGIDI), machine-learning–based Random Forest, Procrustes analysis, and network-based integration of phenotypic and SSR data were implemented using R and Python. Analyses and visualizations were performed with R packages (*stats*, *factoextra*, *cluster*, *GGally*, *ggfortify*) and Python libraries (*pandas*, *numpy*, *scikit-learn*, *networkx*, *matplotlib*, *seaborn*, *scipy*). Statistical significance was assessed at *p* < 0.05 unless otherwise stated.

## Results

### Variability among Mutants

Seventeen traits in the mutants exhibited significant genotypic effects according to ANOVA (S2 Table). The largest genotypic variance was observed for grain yield hill^-1^ (278.22), while the highest phenotypic variance was recorded for filled grains panicle^-1^ (446.24) (Table 1). Grain yield hill^-1^ expressed the highest genotypic coefficient of variation (29.07), whereas sterile grains panicle^-1^ showed the maximum phenotypic coefficient of variation (61.20). Broad-sense heritability estimates ranged from medium to high for days to flowering, days to maturity, plant height, total tillers hill^-1^, effective tillers hill^-1^, panicle length, flag leaf length, filled grains panicle^-1^, grain length, grain breadth, 1000-grain weight, grain yield hill^-1^, straw yield hill^-1^, and harvest index. Genetic advance as percentage of mean was lowest for days to maturity (1.92%) and highest for grain yield hill⁻¹ (59.02%).

**Table 1.**
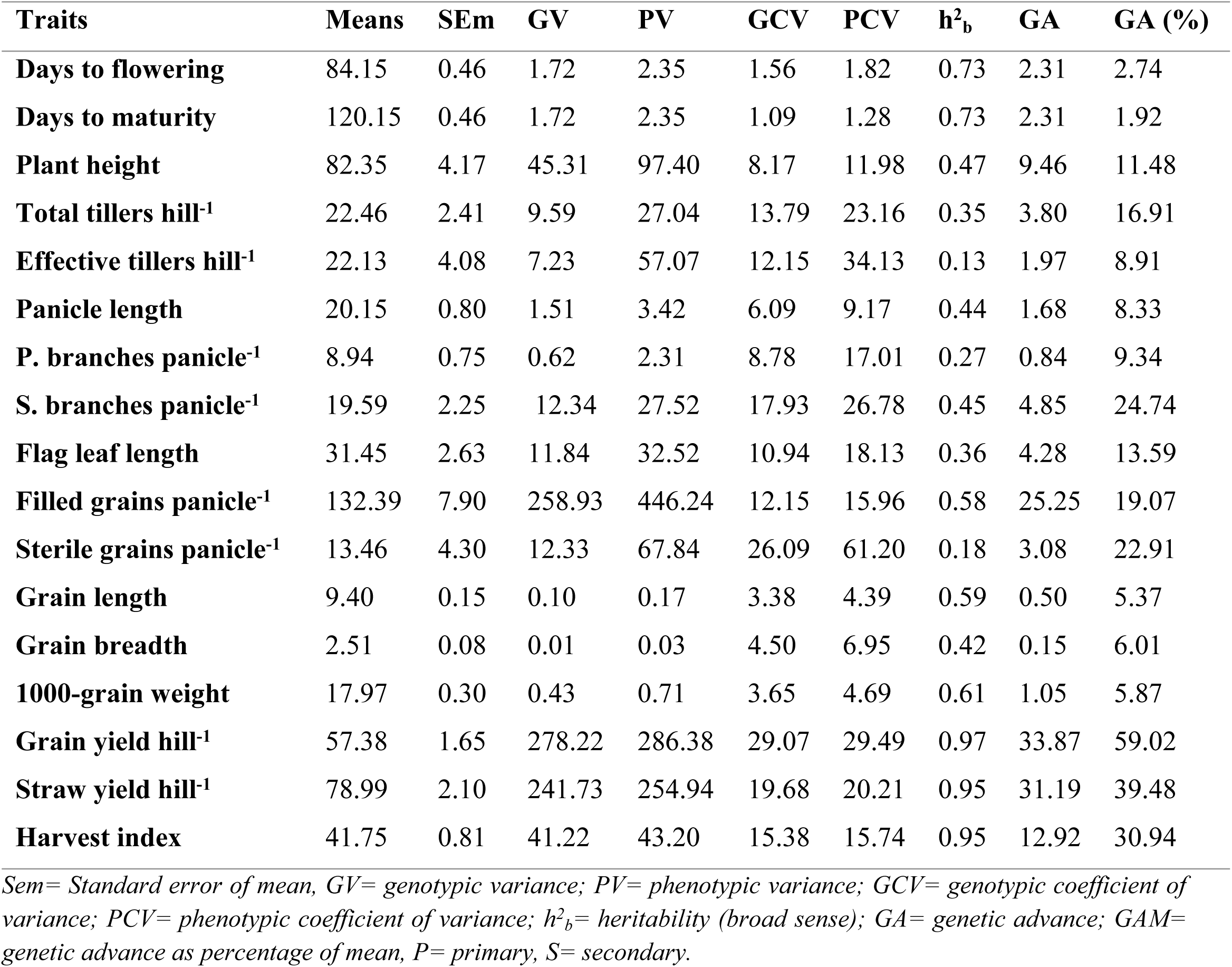
Estimation of genetic parameters in 17 characters of 100 mutants (M3) and their parent.

| Traits | Means | SEm | GV | PV | GCV | PCV | $h^2_b$ | GA | GA (%) |
| --- | --- | --- | --- | --- | --- | --- | --- | --- | --- |
| Days to flowering | 84.15 | 0.46 | 1.72 | 2.35 | 1.56 | 1.82 | 0.73 | 2.31 | 2.74 |
| Days to maturity | 120.15 | 0.46 | 1.72 | 2.35 | 1.09 | 1.28 | 0.73 | 2.31 | 1.92 |
| Plant height | 82.35 | 4.17 | 45.31 | 97.40 | 8.17 | 11.98 | 0.47 | 9.46 | 11.48 |
| Total tillers hill<sup>-1</sup> | 22.46 | 2.41 | 9.59 | 27.04 | 13.79 | 23.16 | 0.35 | 3.80 | 16.91 |
| Effective tillers hill<sup>-1</sup> | 22.13 | 4.08 | 7.23 | 57.07 | 12.15 | 34.13 | 0.13 | 1.97 | 8.91 |
| Panicle length | 20.15 | 0.80 | 1.51 | 3.42 | 6.09 | 9.17 | 0.44 | 1.68 | 8.33 |
| P. branches panicle<sup>-1</sup> | 8.94 | 0.75 | 0.62 | 2.31 | 8.78 | 17.01 | 0.27 | 0.84 | 9.34 |
| S. branches panicle<sup>-1</sup> | 19.59 | 2.25 | 12.34 | 27.52 | 17.93 | 26.78 | 0.45 | 4.85 | 24.74 |
| Flag leaf length | 31.45 | 2.63 | 11.84 | 32.52 | 10.94 | 18.13 | 0.36 | 4.28 | 13.59 |
| Filled grains panicle<sup>-1</sup> | 132.39 | 7.90 | 258.93 | 446.24 | 12.15 | 15.96 | 0.58 | 25.25 | 19.07 |
| Sterile grains panicle<sup>-1</sup> | 13.46 | 4.30 | 12.33 | 67.84 | 26.09 | 61.20 | 0.18 | 3.08 | 22.91 |
| Grain length | 9.40 | 0.15 | 0.10 | 0.17 | 3.38 | 4.39 | 0.59 | 0.50 | 5.37 |
| Grain breadth | 2.51 | 0.08 | 0.01 | 0.03 | 4.50 | 6.95 | 0.42 | 0.15 | 6.01 |
| 1000-grain weight | 17.97 | 0.30 | 0.43 | 0.71 | 3.65 | 4.69 | 0.61 | 1.05 | 5.87 |
| Grain yield hill<sup>-1</sup> | 57.38 | 1.65 | 278.22 | 286.38 | 29.07 | 29.49 | 0.97 | 33.87 | 59.02 |
| Straw yield hill<sup>-1</sup> | 78.99 | 2.10 | 241.73 | 254.94 | 19.68 | 20.21 | 0.95 | 31.19 | 39.48 |
| Harvest index | 41.75 | 0.81 | 41.22 | 43.20 | 15.38 | 15.74 | 0.95 | 12.92 | 30.94 |

*Sem*= Standard error of mean, *GV*= genotypic variance; *PV*= phenotypic variance; *GCV*= genotypic coefficient of variance; *PCV*= phenotypic coefficient of variance; $h^2_b$ = heritability (broad sense); *GA*= genetic advance; *GAM*= genetic advance as percentage of mean, *P*= primary, *S*= secondary.

### Principal Component Analysis (PCA)

Principal component analysis (PCA) of 17 traits in 100 M3 mutants and their parent is presented in Fig 1. The first two principal components (PC1 and PC2) explained 34.94% and 17.18% of the total variation, respectively, accounting for 52.12% of the cumulative variation. The biplot shows the distribution of genotypes (green points) relative to the trait vectors (black arrows). PC1 was primarily associated with plant height, panicle length, primary and secondary branches panicle^-1^, filled grains panicle^-1^, grain yield hill^-1^, and straw yield hill^-1^, indicating that these traits contributed most to the separation of genotypes along this axis. PC2 was influenced by traits such as effective tillers hill^-1^, total tillers hill^-1^, and harvest index. The spread of the points across quadrants reflects the genetic diversity among the mutants, with some genotypes clustering near the parent and others showing distinct trait combinations.

**Fig 1.**
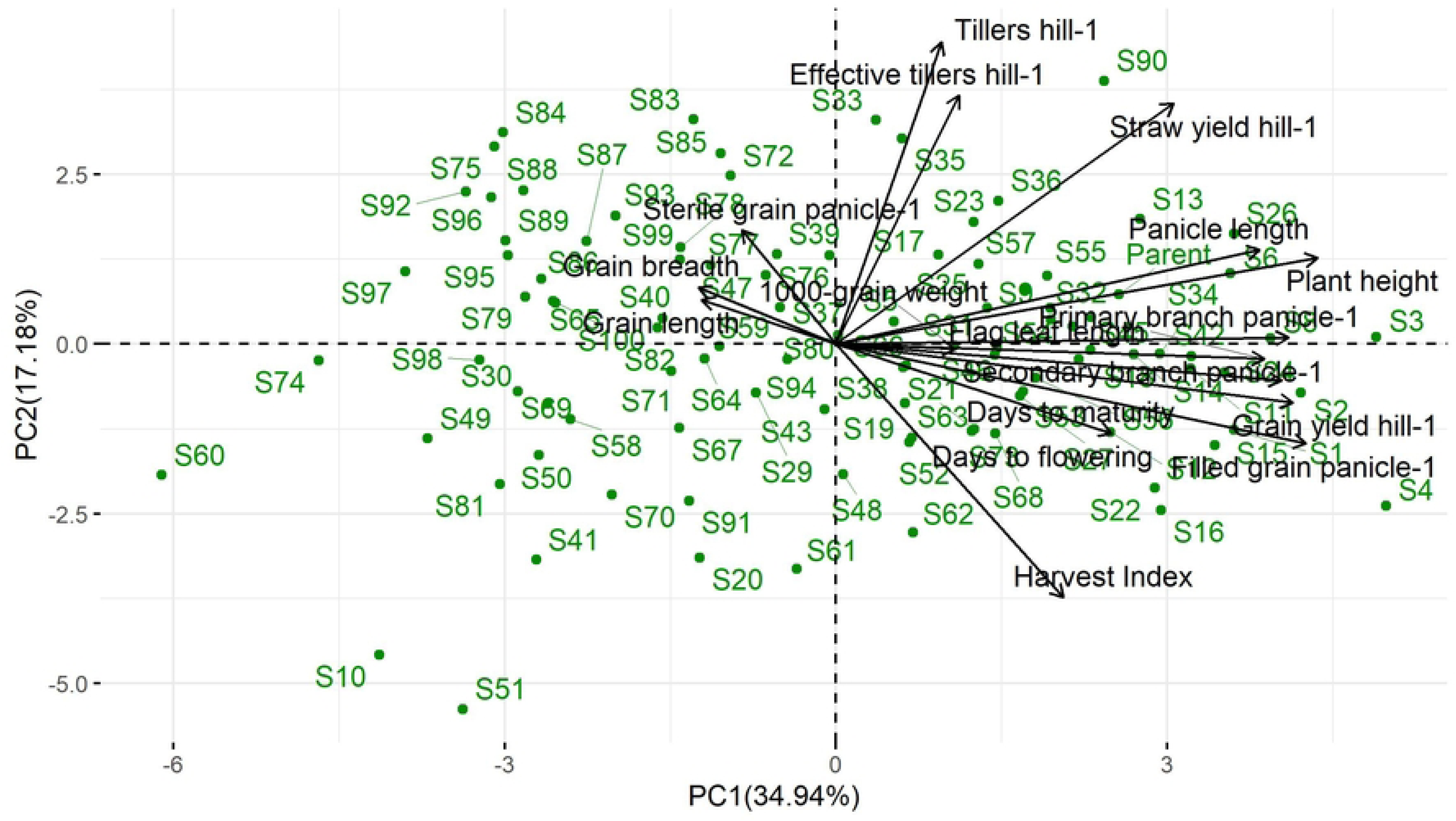
Biplot of principal component analysis (PCA) based on 17 quantitative characteristics, where each point represents the mean of three replicates per genotype; black arrows indicate the standardized active variables contributing to PCA formation.

### Phenotypic correlation among Harvest Index–associated traits

HI was positively and significantly correlated with GYP indicating that allocation of assimilates in the form of increased to grain directly affects HI(Fig 2). In contrast, there was a marked negative correlation between HI and SYP, indicating a biological trade-off between grain filling and vegetative biomass accumulation. This negative correlation reinforces the advantageous effects of balanced biomass partitioning on yield efficiency. Grain yield per plant was significantly and positively correlated with those of the subtraits except grain width, suggesting that a favorable combination of plant architecture and panicle traits can enhance their synergistic effect on grain yield. The panicle length and plant height were also significantly and positively correlated to each other, suggesting that the mutant population displayed corresponding increases in size of vegetative and reproductive organs. Thousand grain weight showed weaker but significant relationships with both grain yield and HI, indicating that grain weight make a separable contribution since it is not the sole determinant of HI. Filled grain number was positively correlated with both grain yield and panicle length, indicating that it is a key component trait for yield.

**Fig 2.**
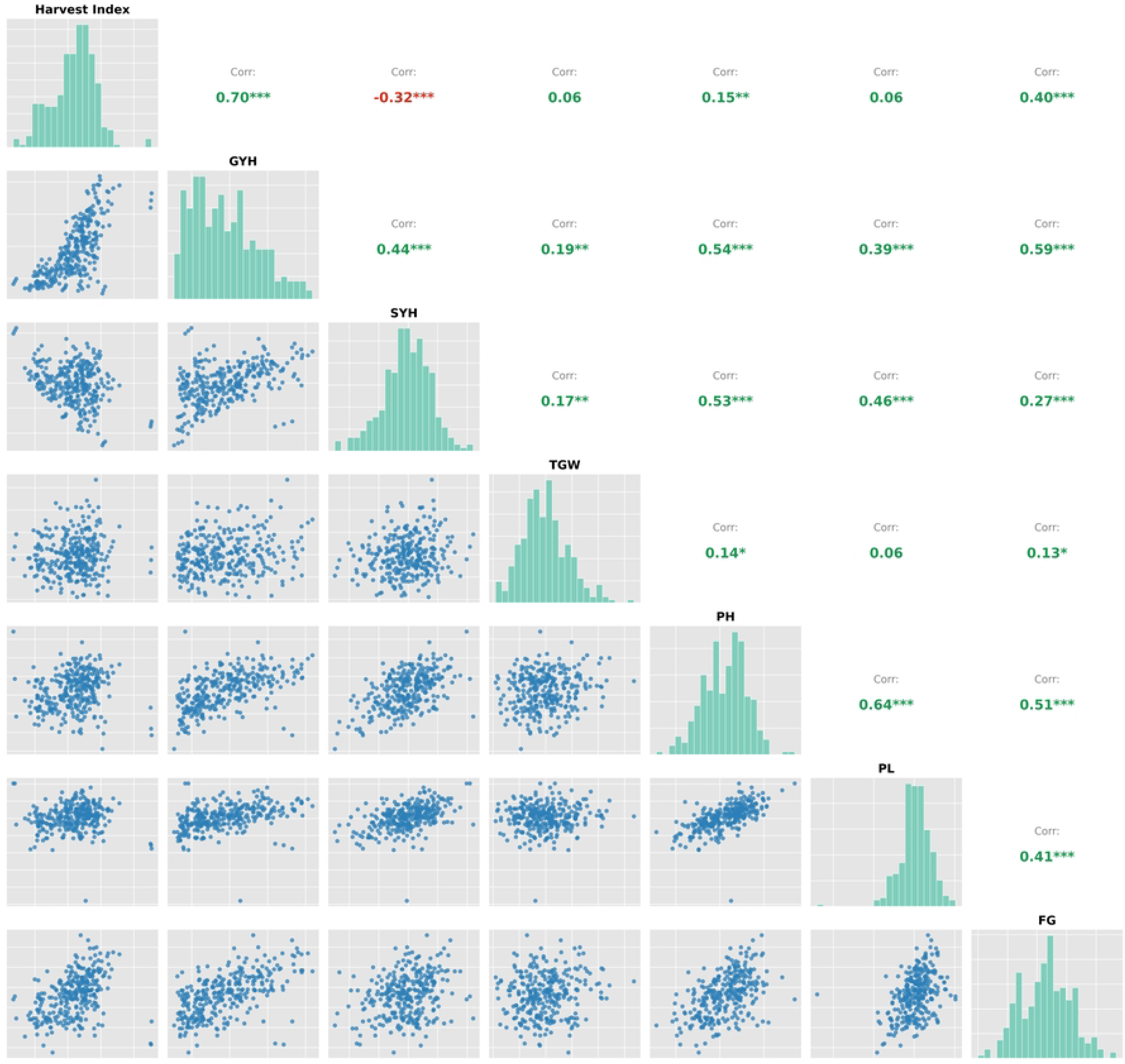
**Phenotypic correlation among seven Harvest Index–associated traits**. Pearson’s correlation matrix showing trait distributions along the diagonal, pairwise scatter plots in the lower triangle, and correlation coefficients with significance levels in the upper triangle. Green and red values represent positive and negative correlations, respectively. Asterisks indicate significance at *p* < 0.05*, p < 0.01, and p < 0.001*.

### Integrated multi-trait selection and prediction of harvest index

MGIDI analysis revealed pronounced multi-trait variability among genotypes, with ten mutants (S-26, S-36, S-83, S-76, S-42, S-75, S-33, S-28, S-3, and S-11) exhibiting the lowest MGIDI values and therefore the closest proximity to the ideotype, while the parental genotype showed inferior overall multi-trait performance (Fig. 3a). The superiority of MGIDI-selected mutants was visually corroborated using a Z-score–standardized heatmap, which revealed consistently favorable profiles for grain yield, HI, filled grains per panicle, and thousand-grain weight in elite mutants compared with the parent. Phenological traits were also clearly separated from yield-related ones at the second level of partitioning by hierarchical clustering and hence were showing a partially independent control which supports the biological significance of the multi-trait selection (Fig. 3b; S3 Table). Using these patterns as a base, it is also observed from the PLS regression that the relationship between predicted and measured HI values was highly linear with tightly clustered data points situated close to the 1:1-line of reference across all populations which clearly indicates that HI can be accurately predicted from such multi-trait integrated dataset (Fig. 3c). To dissect the most important factors for this prediction, Random Forest analysis showed that grain yield per hill and straw yield per hill strongly ruled HI determination; other morpho-physiological characters played a marginal role once biomass partitioning was considered (Fig. 3d).

**Fig 3.**
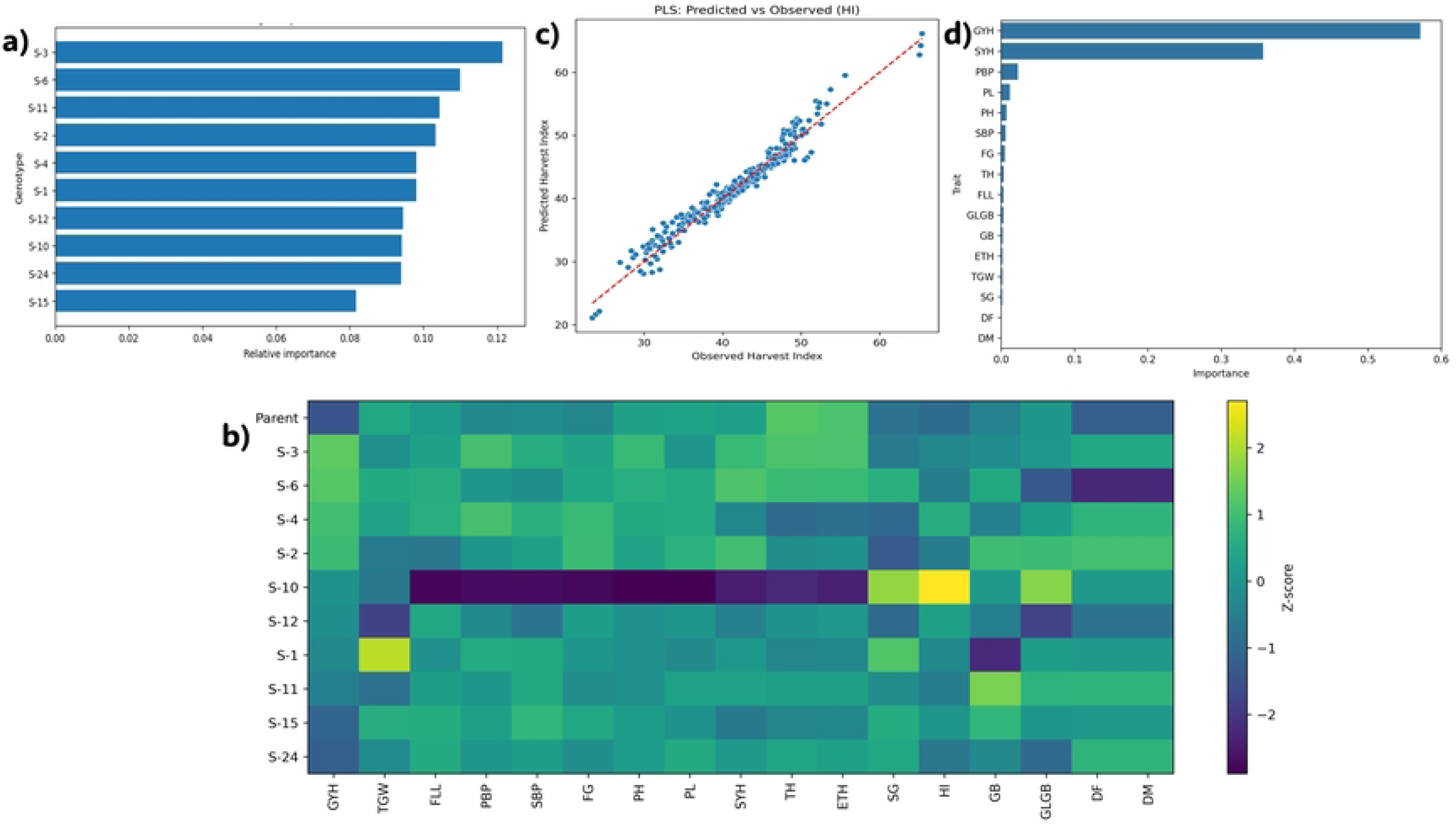
Integrated multi-trait selection and prediction of harvest index. MGIDI analysis identifies elite mutants with superior combined agronomic performance (a) which is corroborated by Z-score standardized heatmap visualization of trait profiles (b) Partial Least Squares regression confirms accurate multivariate prediction of harvest index (c) while Random Forest feature-importance analysis highlights biomass production and partitioning traits as the primary drivers of harvest index variation (d).

### Trait-wise variability among MGIDI-selected elite mutants

The RF feature-importance for six yield-contributing traits among the top 10 mutant genotypes chosen by MGIDI index is given in Fig. 4 using combined violin–box plots. This violin width is proportional to the observation density and includes embedded boxplots that indicate median and interquartile range, while jittered points correspond to replication-level values. Grain yield hill⁻¹ (Fig. 4a) considerably varied among mutants, whereas some genotypes developed with higher values of yield and narrow distributions, indicating stable yield performance over three replications. Straw yield hill⁻¹ (Fig. 4b) differed substantially, which indicated that biomass production and distribution varied by genotype. For the architectural features of the panicle, panicle length (Fig. 4d) displayed low divergence, and the numbers of primary branches panicle⁻¹ (Fig. 4c) showed relatively low dispersion across most genotypes. In contrast, secondary branches panicle⁻¹ (Fig. 4e) exhibited pronounced differences among mutants, characterized by wider distribution profiles in selected lines, implying that secondary branching is a major source of variation within the elite group. Similarly, filled grains panicle⁻¹ (Fig. 4f) indicated significant variation among the mutants, in that the best lines consistently showed high and stable filled grain number suggesting its role in improved GY.

**Fig 4.**
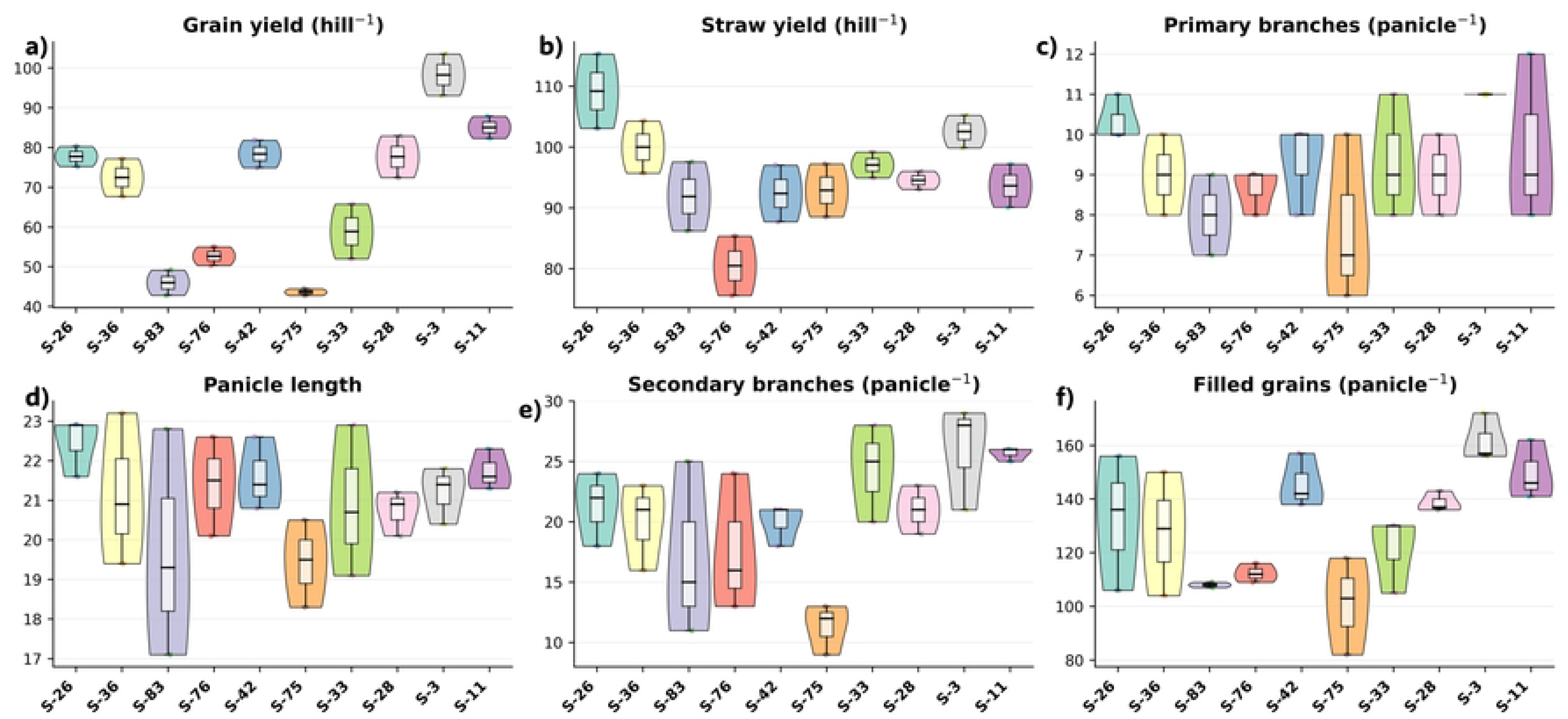
Distribution of key yield-contributing traits among MGIDI-selected elite mutant genotypes. Combined violin–box plots illustrate the replication-level distribution, central tendency, and variability of six yield-related traits across the top 10 elite mutants (S-26, S-36, S-83, S-76, S-42, S-75, S-33, S-28, S-3, and S-11) selected by the MGIDI index: (a) grain yield hill⁻¹, (b) straw yield hill⁻¹, (c) primary branches panicle⁻¹, (d) panicle length, (e) secondary branches panicle⁻¹, and (f) filled grains panicle⁻¹. The violin width represents data density, embedded boxplots indicate median and interquartile range, and points denote individual replicates.

### Association of Grain Yield with Yield-Contributing Traits

The strongest correlation with grain yield was observed with FGPP (r = 0.59, p < 0.001; Fig. 5a). Traits of panicle architecture showed a moderate and extremely significant correlation with grain yield. Panicle length was positively correlated with yield (r = 0.39, p = 1.41 × 10⁻¹²; Fig. 5b), indicating that more spikelet-bearing units of longer panicles are favourable to enhance the yield potential. Also, at primary (r = 0.38, p = 3.91 × 10⁻¹²; Fig. 5c) and number of secondary branches per panicle (r = 0.44, p = 1.55 × 10⁻¹⁵; Fig. 5d) varied significantly and had positive correlations with yield, emphasizing the role of panicle branching in sink size/dry grain formation. Straw yield per hill was also significantly positively correlated with grain yield (r = 0.44, p = 8.88 × 10⁻¹⁶; Fig. 5e), suggesting that strong vegetative growth can be translated into enhanced reproductive output, which represents the effect of plant vigor on yield production. On the other hand, 1000-grain weight showed a weaker but positive and significant correlation with grain yield (r = 0.19, p = 0.001; Fig. 5f), indicating that grain size is likely to have a relatively minor effect on yield variation compared to those related to panicle structure and grain number.

**Figure 5.**
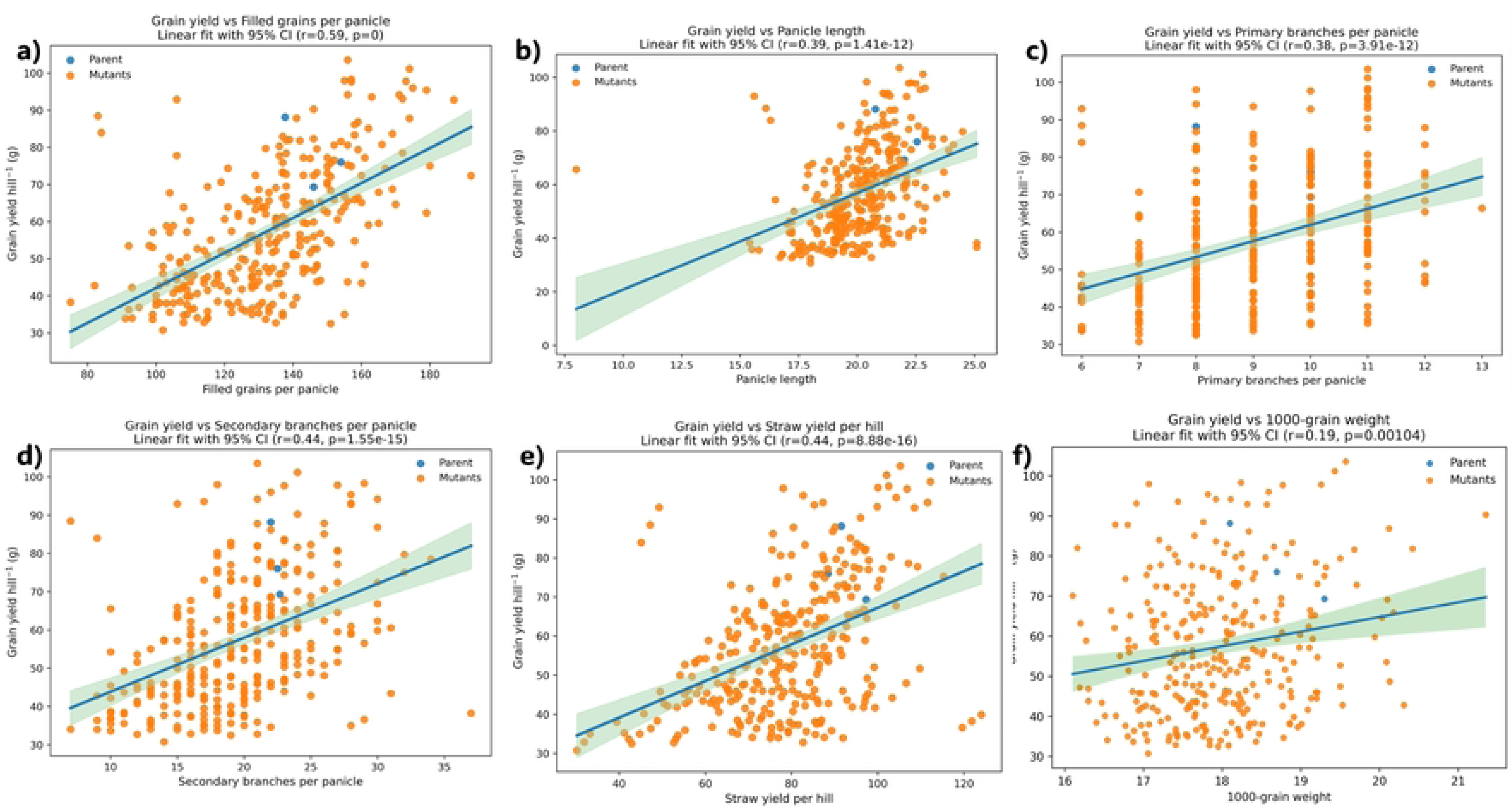
Relationship between grain yield and yield-contributing traits. Linear regression analyses (±95% CI) illustrate associations between grain yield per hill and (a) filled grains per panicle, (b) panicle length, (c) primary branches per panicle, (d) secondary branches per panicle, (e) straw yield per hill, and (f) 1000-grain weight, with filled grains per panicle and biomass-related traits showing the strongest effects.

### Genetic Diversity and Differentiation in Rice Mutants

Thirty SSR markers were analyzed among rice mutants, and their high genetic diversity and population structure were observed. Major Allele Frequency (MAF) ranged from 0.48 (RM124) to 0.96 (RM133), averaging 0.76; low MAF (≤0.55) was observed in RM124, RM215, and RM431, while high MAF (≥0.90) occurred in RM133, RM171, RM237, RM433, and RM484. Gene diversity (He) ranged from 0.0768 (RM152) to 0.4968 (RM237), with a mean value of approximately 0.315. The highest gene diversity was observed in markers RM237 (0.4968), RM408 (0.4950), RM433 (0.4950), RM447 (0.4950), RM495 (0.4872), and RM514 (0.4712). PIC values ranged from 0.12 (RM215) to 0.37 (RM237, RM408) and were higher in the markers RM237, RM408, RM484 and RM433. The highest Ho was observed in RM44 and RM152 (0.92 and 0.96), whereas He was evenly dispersed among markers such as RM125 (0.4118) (Table 2).

**Table 2.**
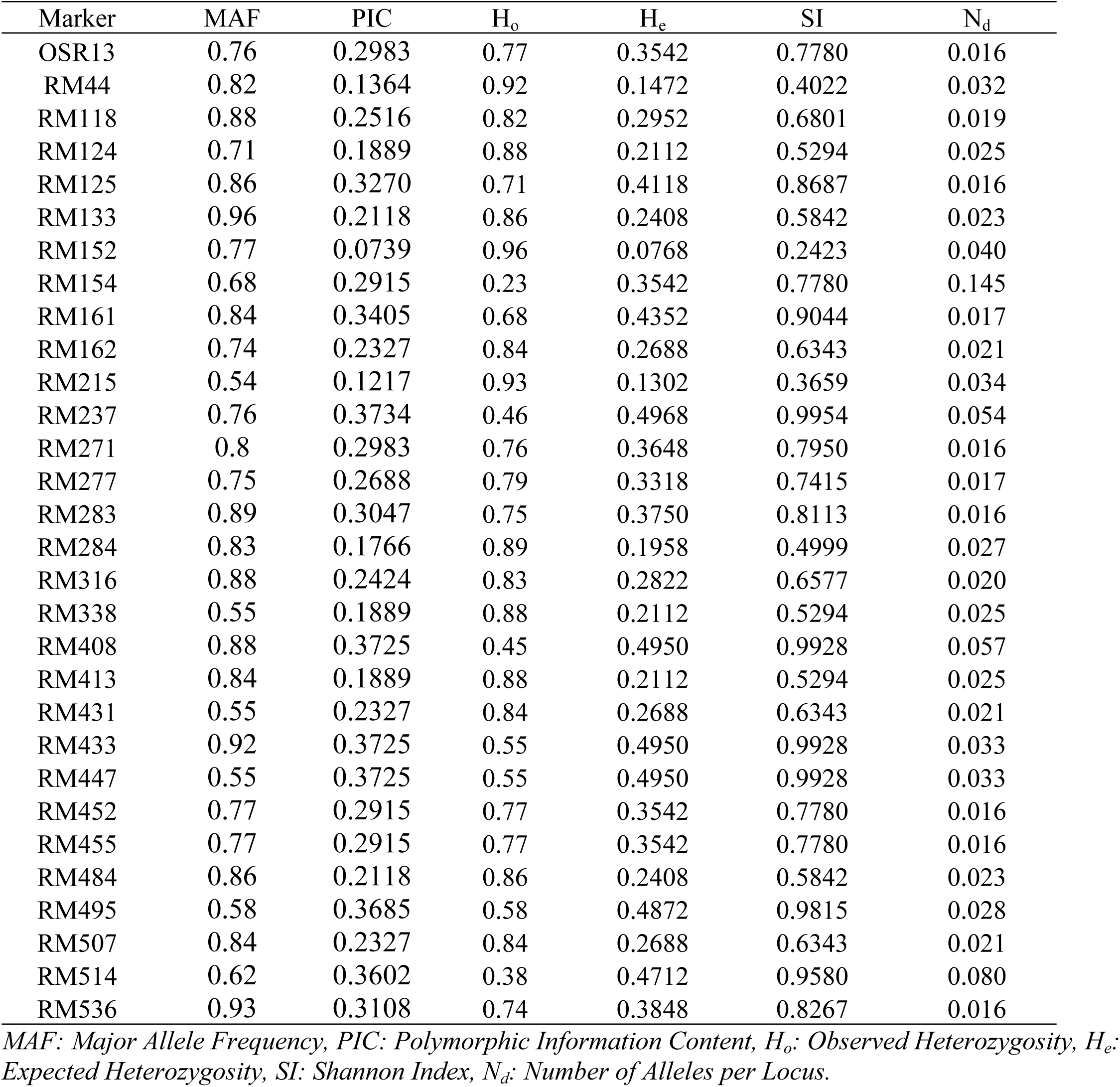
Summary of genetic diversity parameters for 30 SSR markers used in 100 SA-induced rice mutants.

| Marker | MAF | PIC | H<sub>o</sub> | H<sub>e</sub> | SI | N<sub>d</sub> |
| --- | --- | --- | --- | --- | --- | --- |
| OSR13 | 0.76 | 0.2983 | 0.77 | 0.3542 | 0.7780 | 0.016 |
| RM44 | 0.82 | 0.1364 | 0.92 | 0.1472 | 0.4022 | 0.032 |
| RM118 | 0.88 | 0.2516 | 0.82 | 0.2952 | 0.6801 | 0.019 |
| RM124 | 0.71 | 0.1889 | 0.88 | 0.2112 | 0.5294 | 0.025 |
| RM125 | 0.86 | 0.3270 | 0.71 | 0.4118 | 0.8687 | 0.016 |
| RM133 | 0.96 | 0.2118 | 0.86 | 0.2408 | 0.5842 | 0.023 |
| RM152 | 0.77 | 0.0739 | 0.96 | 0.0768 | 0.2423 | 0.040 |
| RM154 | 0.68 | 0.2915 | 0.23 | 0.3542 | 0.7780 | 0.145 |
| RM161 | 0.84 | 0.3405 | 0.68 | 0.4352 | 0.9044 | 0.017 |
| RM162 | 0.74 | 0.2327 | 0.84 | 0.2688 | 0.6343 | 0.021 |
| RM215 | 0.54 | 0.1217 | 0.93 | 0.1302 | 0.3659 | 0.034 |
| RM237 | 0.76 | 0.3734 | 0.46 | 0.4968 | 0.9954 | 0.054 |
| RM271 | 0.8 | 0.2983 | 0.76 | 0.3648 | 0.7950 | 0.016 |
| RM277 | 0.75 | 0.2688 | 0.79 | 0.3318 | 0.7415 | 0.017 |
| RM283 | 0.89 | 0.3047 | 0.75 | 0.3750 | 0.8113 | 0.016 |
| RM284 | 0.83 | 0.1766 | 0.89 | 0.1958 | 0.4999 | 0.027 |
| RM316 | 0.88 | 0.2424 | 0.83 | 0.2822 | 0.6577 | 0.020 |
| RM338 | 0.55 | 0.1889 | 0.88 | 0.2112 | 0.5294 | 0.025 |
| RM408 | 0.88 | 0.3725 | 0.45 | 0.4950 | 0.9928 | 0.057 |
| RM413 | 0.84 | 0.1889 | 0.88 | 0.2112 | 0.5294 | 0.025 |
| RM431 | 0.55 | 0.2327 | 0.84 | 0.2688 | 0.6343 | 0.021 |
| RM433 | 0.92 | 0.3725 | 0.55 | 0.4950 | 0.9928 | 0.033 |
| RM447 | 0.55 | 0.3725 | 0.55 | 0.4950 | 0.9928 | 0.033 |
| RM452 | 0.77 | 0.2915 | 0.77 | 0.3542 | 0.7780 | 0.016 |
| RM455 | 0.77 | 0.2915 | 0.77 | 0.3542 | 0.7780 | 0.016 |
| RM484 | 0.86 | 0.2118 | 0.86 | 0.2408 | 0.5842 | 0.023 |
| RM495 | 0.58 | 0.3685 | 0.58 | 0.4872 | 0.9815 | 0.028 |
| RM507 | 0.84 | 0.2327 | 0.84 | 0.2688 | 0.6343 | 0.021 |
| RM514 | 0.62 | 0.3602 | 0.38 | 0.4712 | 0.9580 | 0.080 |
| RM536 | 0.93 | 0.3108 | 0.74 | 0.3848 | 0.8267 | 0.016 |

MAF: Major Allele Frequency, PIC: Polymorphic Information Content, H<sub>o</sub>: Observed Heterozygosity, H<sub>e</sub>: Expected Heterozygosity, SI: Shannon Index, N<sub>d</sub>: Number of Alleles per Locus.

### Cluster Analysis of Rice Mutants

Agglomerative hierarchical clustering of 100 SA-induced rice mutants based on 30 SSR markers revealed a clear multi-level dendrogram shown in Fig 6. A linkage distance of ∼15 separated three major clusters. Cluster A (∼83 mutants) exhibited moderate heterogeneity, with sub-branches merging between distances 8–14. Cluster B (∼12 mutants, e.g., SA78, SA95, SA87, SA89, SA93, SA99, SA98, SA94, SA88, SA90, SA32, SA84) was compact, coalescing by ∼12, indicating higher within-group similarity. Cluster C (5 mutants; SA80, SA81, SA83, SA86, SA100) was the most homogeneous, with internal joins complete by ∼9–10. Clusters B and C were closer to each other than to Cluster A, merging at ∼16–17, while their union with Cluster A occurred at ∼24–25, showing pronounced separation of the majority lineage. Short terminal branches (<3–4) revealed near-identical pairs, notably SA81–SA83, SA90–SA32, and SA95–SA87, with no persistent single-sample outliers.

**Fig 6.**
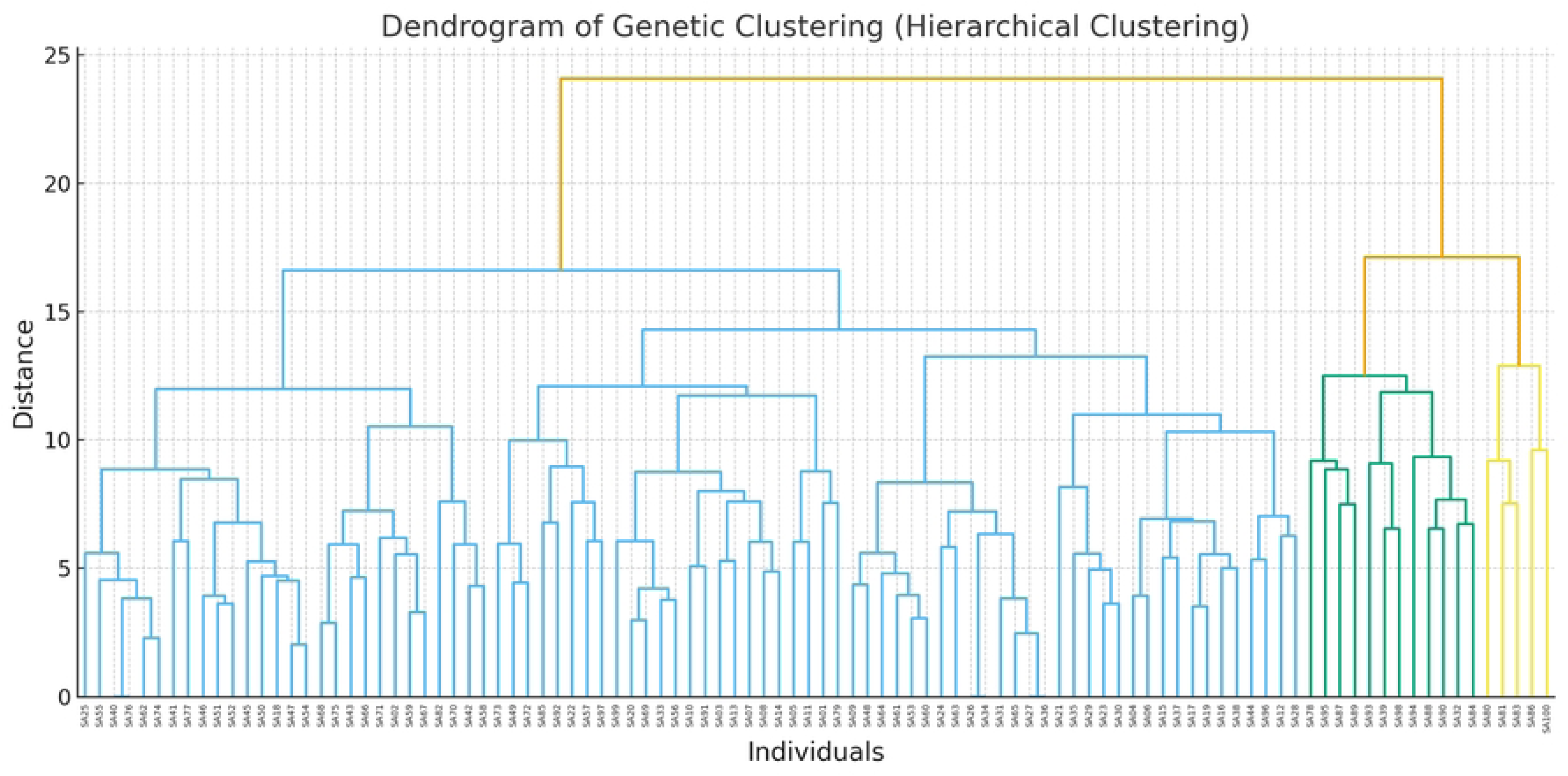
Dendrogram of 100 SA-induced rice mutants based on 30 SSR markers (agglomerative hierarchical clustering)

### Population Structure Analysis

The population structure of 100 SA-induced rice mutants was assessed using STRUCTURE v2.3.4, employing data from 30 SSR markers. According to Evanno’s ΔK method, the analysis revealed two well-defined subgroups (Fig. 7a), which were further subdivided into “pure” and “admixture” categories based on the proportion of genetic assignment. Genotypes with an assignment proportion greater than 0.8 were classified as “pure,” while those with an assignment proportion less than 0.8 were designated as “admixtures. “Population I demonstrated a strong membership to Cluster 1, whereas Population II showed a predominant membership to Cluster 2 (Fig. 7b). Notably, some genotypes displayed flexible memberships, indicating admixture, with proportions below 0.8. The genetic divergence between the two populations, measured by the average allelic frequency (Fst), was found to be 0.0437, indicating low genetic differentiation. Further supporting this genetic distinction, the average alpha value was calculated to be 0.4189, and Fst values for Cluster 1 and Cluster 2 were 0.0026 and 0.3584, respectively, confirming distinct genetic structuring between the populations. In terms of model parameters, the log(alpha) value (Fig. 7c) stabilized after initial fluctuations, signifying the system’s convergence. The Fst_2 parameter (Fig. 7d) exhibited moderate variation and approached a plateau, while the Ln Likelihood (Fig. 7e) demonstrated convergence post the burn-in phase, further supporting the robustness of the population structure and genetic differentiation observed between the two groups.

**Fig 7.**
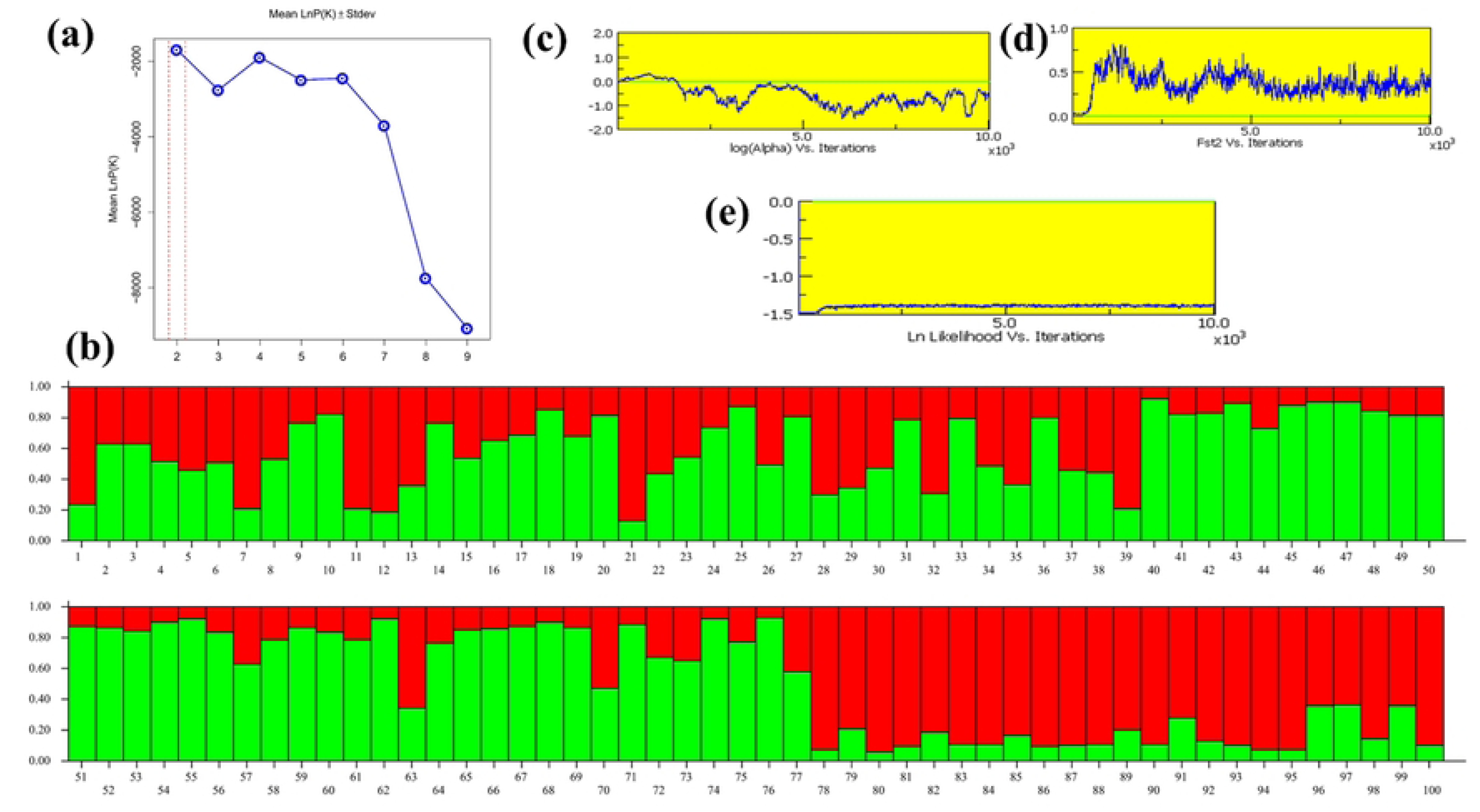
Population structure analysis of SA-induced rice mutants. (a) LnP(D) and ΔK values used to determine the optimal number of genetic clusters. (b) Structure bar plot at K = 2 showing the genetic composition of 100 SA-induced rice mutants based on 30 SSR markers. Logarithmic model-parameter plots across iterations: (c) log (Alpha) vs. iterations, (d) Fst2 vs. iterations, and (e) Ln Likelihood vs. iterations.

### Genetic Relationships among Mutant Genotypes

Pairwise comparison of mutants based on genetic distance revealed varying levels of similarity and divergence (Fig 8, Table 3). Mutant pairs such as SA26 vs. SA34, SA27 vs. SA36, and SA40 vs. SA76 exhibited a genetic distance of 0.000, indicating they are genetically identical and possibly represent duplicates or replicates. Similarly, SA27 vs. SA65 and SA36 vs. SA65 showed very low genetic distances (0.033), suggesting extremely close relationships with a high degree of allele sharing. In contrast, some mutant pairs displayed substantial divergence. SA92 vs. SA93 showed a genetic distance of 0.667, representing a genetically distinct branch within the population. Even greater divergence was observed for SA67 vs. SA80, SA79 vs. SA80, and SA81 vs. SA88, all with 0.700, indicating significant separation from the main population. The highest genetic distance (0.733) was observed between SA81 and SA84, suggesting that these two mutants belong to distinct lineages with little likelihood of recent common ancestry.

**Fig 8.**
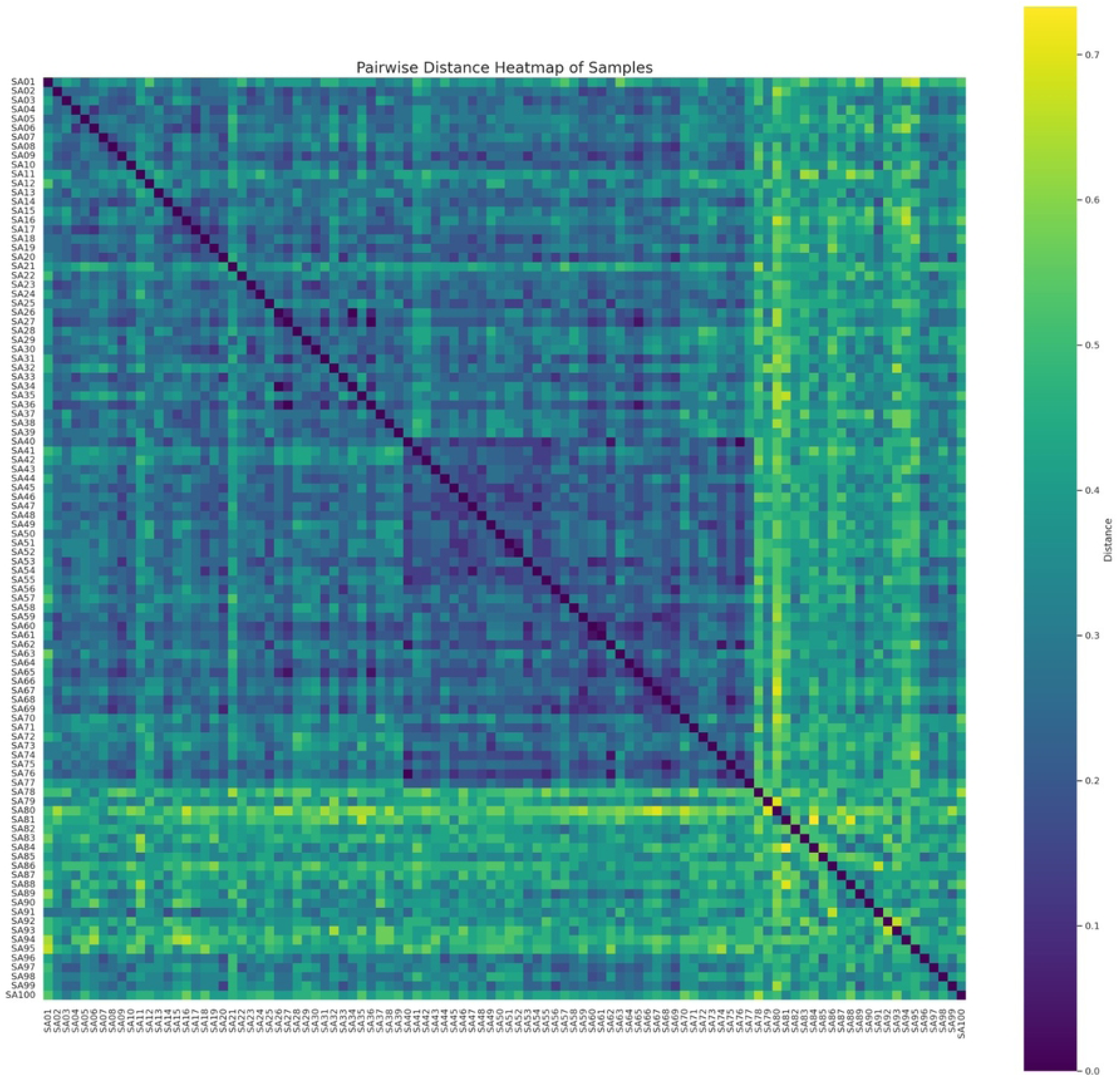
Heatmap of Pairwise Genetic Distances Among Mutant Genotypes

**Table 3.** Annotated Summary of Genetically Significant Relationships Among Mutant Genotypes.

| Mutant Compared | Distance | Relationship Type | Remark |
| --- | --- | --- | --- |
| SA26 vs SA34, SA27 vs SA36, SA40 vs SA76 | 0 | Possibly duplicates | Identical genotypes; likely replicates |
| SA27 vs SA65, SA36 vs SA65, | 0.033 | Shared alleles, high similarity | Very close relationship; high allele similarity |
| SA92 vs SA93 | 0.667 | Genetically distinct branch | Significant divergence from common population |
| SA67 vs SA80, SA79 vs SA80, SA81 vs SA88 | 0.7 | Genetically distinct branch | Significant divergence from common population |
| SA81 vs SA84 | 0.733 | Likely different lineage | Highly differentiated, unlikely recent ancestry |

### Genetic diversity and differentiation among SA-induced rice genotypes

The ordination aligned well with a large and unclosed spread of genotypes observed in the two-dimensional space, suggesting high genetic heterogeneity within the mutant population. The SSR ordination (orange points) revealed a large spatial separation of genotypes, indicative of much allelic variation at loci for the SSRs (Fig 9a; S4 Table). Genotypes were not confined to a single cluster but rather occupied distinct positions throughout the ordination space, suggesting the presence of diverse genetic backgrounds generated through sodium azide mutagenesis. This effect was even more obvious in the overlay of phenotypic ordination (blue symbols), which also revealed that, within mutants, several genotypes are located at isotopically extreme positions along both aligned dimensions (Fig 9b; S4 Table). Several mutants (e.g., S-3, S-4, S-6, S-10, S-11 and S12, and isolate 15) were positioned at a distance from the central cluster indicating that these genotypes are genetically distinct in comparison with many other genotypes.

**Fig 9.**
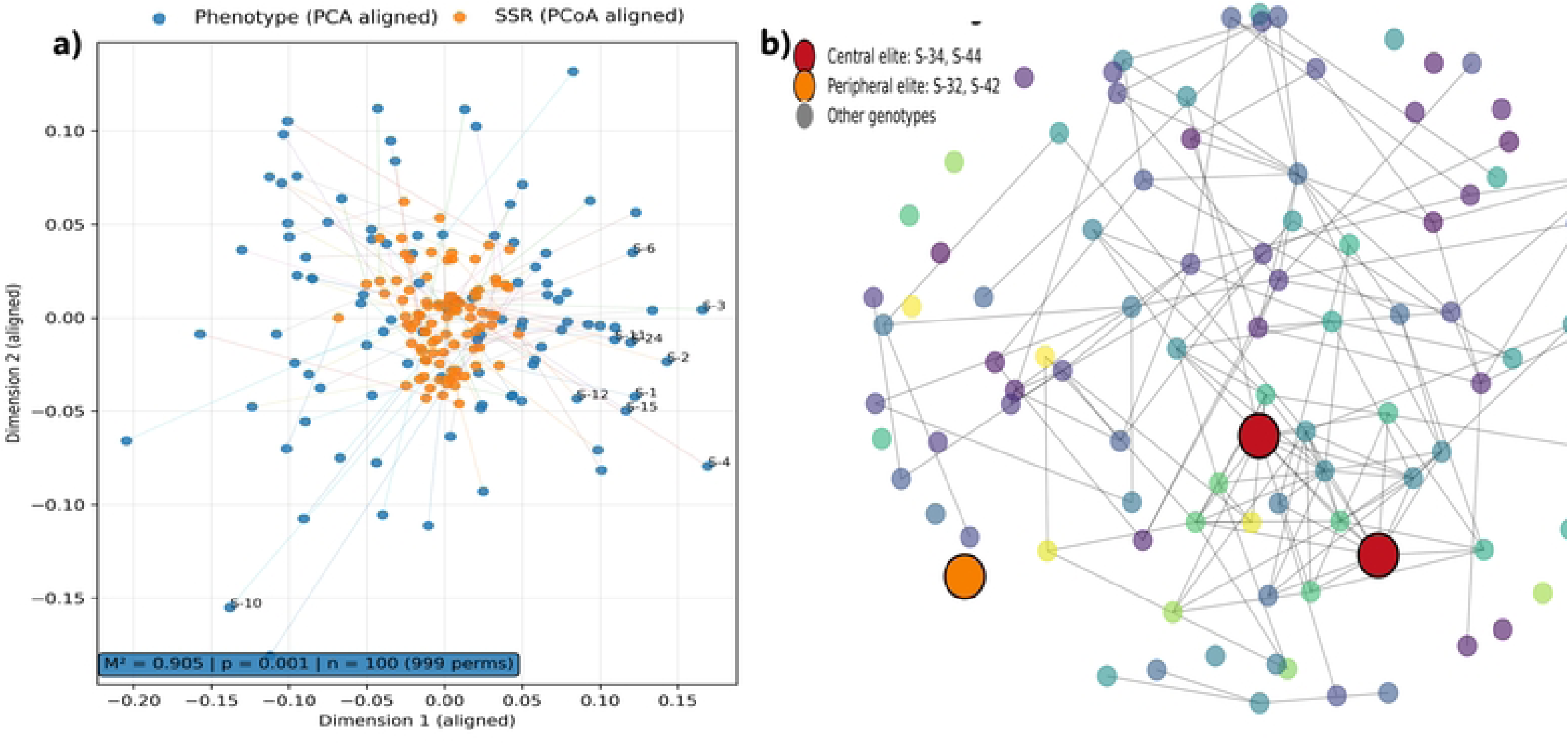
Genetic diversity and differentiation among SA-induced rice genotypes. (a) Procrustes-aligned phenotypic (PCA) and SSR-based genetic (PCoA) ordinations showing broad genotype dispersion and concordance between molecular and phenotypic variation. (b) Network visualization of genetic relationships, highlighting central and peripheral elite genotypes and illustrating extensive genetic differentiation within the mutant population.

## Discussion

The integration of sodium azide mutagenesis with multi-trait selection indices and machine learning provides a comprehensive framework for rice improvement. This study demonstrates that sodium azide generates substantial phenotypic and genetic diversity that can be efficiently exploited using modern analytical tools to identify elite genotypes with superior harvest index (HI) and yield potential. The combined use of morphological evaluation, SSR-based diversity analysis, and multivariate prediction models offers insight into the genetic architecture of HI and supports germplasm utilization in breeding programs. As a potent point mutagen, sodium azide induces mutations ranging from silent substitutions to functionally significant DNA changes mediated through O-acetylserine sulfhydrylase-generated metabolites (Gruszka et al., 2012; Viana et al., 2019).

### Phenotypic Variability and Principal Component Analysis

The screening of 100 sodium azide-induced mutants for seventeen agronomic traits showed the wide variation in phenotypic spectrum. The highest genotypic variance (278.22) and genotypic coefficient of variation (29.07%) about grain yield hill⁻¹ confirms the successful development of variability in this most important economic trait in the present investigation as also reported by Al Mamun et al. (2024). Principal component analysis described 52.12% of the variance, PC1 (34.94%) was mainly related to yield and biomass traits and PC2 (17.18%) was associated with tiller number and HI, which indicates that genetic control is partially independent. The genotypes were distributed in all PCA quadrants, indicating the efficacy of sodium azide for broadening phenotypic variation and creating diverse trait combinations for breeding (Singh et al., 2013).

### Phenotypic Correlations and Trade-offs in Harvest Index

The significant positive correlation of HI with grain yield in the current study is in agreement with previous reports that HI increase contributes to the yield gain in modern rice (Yang and Zhang, 2010; Gu et al., 2023). On the other hand, the negative correlation of HI with straw yield reflects the biological compromises between vegetative biomass and grain filling (Zhang et al., 2022). The moderate to high positive correlations between grain yield and architectural traits such as plant height, panicle length and filled grains per panicle further demonstrate that sink size (in particular spikelets per panicle) is indeed at the center of yield potential (Hill et al., 2024). The positive correlation of filled grains to yield (r = 0.59, p < 0.001) verifies sink limitation in improving yield. In contrast, the weaker correlation between thousand-grain weight and yield or HI (r = 0.19, p = 0.001) indicates that grain number-related traits contribute more substantially than grain size, consistent with classical yield component theory (Khush, 1995; Yamamoto et al., 2021).

### Multi-Trait Selection Using MGIDI

MGIDI success in identifying top performing elite mutants (S-26, S-36, S-83, S-76, S-42, S-75, S-33, S-28, S-3 and S-11) combines agronomic performance. In comparison with traditional indices MGIDI has the advantages of including trait correlation structure and being calculated as a distance to the ideotype (Olivoto and Nardino, 2021). Recent research shows that it is effective at achieving balanced gains, and it outperforms the Smith-Hazel and FAI-BLUP methods during hemispheric competition (Debnath et al., 2024; Sritharan et al., 2025). MGIDI has achieved selection gains of 1.86% to 75.4% in rice (Pallavi et al., 2024; Al Mamun et al., 2024) depending on heritability and environment. The separation of yield-related and phenological traits in the hierarchical clustering indicates that genetic regulation is at least partially independent from one another, which is consistent with the clustering of distinct QTL as reported previously (Deshmukh et al., 2021). The divergence at elite mutants in traits such as number of secondary branches and number of filled grains per panicle suggests that high-performance mutants are derived from mutagenesis of multiple genetic pathways and thus provides breeding flexibility (Liu et al., 2025).

### Machine Learning Prediction of Harvest Index

HI estimates from both PLS regression and RF analyses complemented each other. Observing an excellent agreement between the predicted and observed HI values further confirms the efficiency of multivariate modeling. The HI was dominated by the predictors of grain yield and straw yield according to the RF, which confirmed that biomass partitioning and source–sink relationships are essential (Yang and Zhang, 2010). While deep learning has shown impressive results in large tabular datasets, recent work has shown that RF and related ensemble models tend to outperform DL for small-to-medium tabular datasets, alongside interpretability (Gill et al., 2022; Liu et al., 2025). The combination of machine learning with quantitative genetics can hasten the pace of breeding because the input of the former may get more powerful when applied to high-throughput phenotyping and genomic selection (Singh et al. 2016; Washburn et al. 2021)

### Genetic Diversity and Population Structure

Analysis of 30 SSR markers revealed substantial genetic variation. MAF ranged from 0.48 to 0.96 (mean = 0.76), with low-MAF markers (RM124, RM215, RM431) showing higher diversity and utility for resolving closely related genotypes (Singh et al., 2024). High-MAF markers (RM133, RM171, RM237, RM433, RM484) among closely related species can be indicative of conserved or recently fixed regions (Salem et al., 2024). The PIC values varied between 0.12 to 0.37, and the gene diversity varied from 0.0678 to 0.4968, the most informative markers were RM237 (0.4968), RM408 (0.4950), RM433 (0.4950), RM447 (0.4950), RM495 (0.4872), and RM514 (0.4712).

STRUCTURE analysis revealed two subgroups (Fst = 0.0437, alpha = 0.4189, Fst₁ = 0.0026, Fst₂ = 0.3584) with moderate differentiation overall but strong divergence between Population II. Hierarchical clustering classified the mutants into three groups, although Cluster C was the most homogeneous (SA80, SA81, SA83, SA86, SA100). The pairwise genetic distances were between 0.000 (SA26 (vs. SA34); SA27 (vs. SA36); SA40 (vs. SA76)) and 0.733 (SA81 (vs. SA84)). For instance, identical or near-isogenic pairs (0–0.033) have likely mutational history, while highly divergent mutants (0.667–0.733) represent novel genetic constitutions, which are potential source of rice gene pool (Wang et al., 2013). There was good agreement between molecular and phenotypic variation based on Procrustes alignment, indicating that these induced mutations were phenotypically consequential (Al Mamun et al. 2024, Wang et al. 2024).

### Integration of Multi-Trait Selection, Machine Learning, and Mutagenesis

This integrated analytical framework leveraging MGIDI-based multi-trait selection, machine learning prediction-based breeding models and genetic diversity assessment used in this study provides a powerful approach for modern crop improvement. This integrative approach directly targets several critical problems of rice breeding: (1) simultaneous gain in multiple correlated traits; (2) unknown relative contribution of trait components to the ideal phenotypes; and (3) insufficient allelic diversity for sustained breeding gain. The agreement between MGIDI selection outcomes and predicted selections by the machine learning approach validates both methods in a complementary sense and increases the confidence that genotypes of high predicted performance are truly ideals. We demonstrate the successful application of MGIDI to sodium azide-induced mutants in this regard by showing that the variations induced can be screened effectively with contemporary selection indices, providing a way to accelerate the selection cycle over what the traditional phenotypic selection approaches would permit (Debnath et al., 2024; Sritharan et al., 2025).

### Implications for Breeding Strategies and Future Directions

The findings from this study have several important implications for rice breeding strategies aimed at improving HI and overall yield potential. The predominance of traits related to biomass production and partitioning in explaining variation in HI, indicates that breeding efforts should target functioning as a system, as opposed to improvement of individual yield components. This suggestion is consistent to a recent trend in breeding, which is emphasizing both biomass production potential and HI in balance for improvement (Gu et al., 2023; Zhang et al., 2022).

Third, the large population of mutants produced via sodium azide mutagenesis allow for the rapid discovery of new alleles that are otherwise not easily accessible in natural germplasm collections. Sodium azide (SA) has succeeded in producing a wide array of broad resistance mutants and benefited economically important traits in rice breeding programs across the globe (Wang et al., 2013; Wang et al., 2024). The chosen elite mutants based on MGIDI analysis are ideal candidates for direct representation in breeding programs as either parental lines or favorable alleles for marker-assisted introgression.

Despite the moderate average PIC value for the SSR markers (0.264), it was of sufficient value for genetic diversity detection and for supporting marker-assisted selection strategies. PIC values from 0.34 to 0.37 indicated that RM237, RM408, RM484 and RM433 are highly informative and very useful for selecting elite genotypes and monitoring genetic diversity in breeding populations. The population structure provides evidence for heterozygosity from inter-population crosses (K = 2, Fst = 0.0437). The high heritability of grain yield, straw yield, and harvest index can provide basis for good selection. The fact that PC1 and PC2 are independent of each other and vary independently suggests that it is indeed possible for trait complexes to be improved at the same time by proper selection, allowing a simultaneous and extensive genetic gain.

There is a need for genomic characterization of elite mutants, multi-environment validation, genomic selection with MGIDI and other algorithms. Understanding the physiological mechanisms driving HI associations will do even more to fine-tune ideotype design. The short-term exploitation of a few high-yielding clusters can be equitably balanced with the long-term utilization of diverging outliers to enable short-term productivity benefits while safeguarding genetic sustainability in rice breeding programs.

## Conclusion

This research demonstrates the combined use of agronomic and molecular analyses as an efficient approach for rice mutant characterization that offers practical guidance for improving mutation breeding programs and effective application of genetic variation in developing elite rice varieties. The integration of multi-trait selection indices, machine learning approaches, and mutagenesis-generated genetic diversity provides valuable resources and tools for future breeding programs aimed at developing high-yielding rice varieties with optimized harvest index and improved adaptation to diverse production environments.

## Acknowledgement

We gratefully acknowledge the Research and Innovation Centre (RIC), Khulna University, for providing financial and institutional support essential for the successful completion of this research.

## Author contributions

**S. M. Abdullah Al Mamun:** Conceptualization, Methodology, Funding acquisition, Project administration, Supervision, Investigation, Writing – original draft: **Md Rezve:** Conceptualization, Investigation, Methodology, Data curation, Formal analysis, Software, Visualization, Writing – original draft, Writing – review and editing**: Md. Borhan Ali Sorker:** Investigation, Methodology, Data curation: **Md. Musabbir Hossain Shoun**: Investigation, Methodology, Data curation**: Mst. Sabiha Sultana:** Validation, Writing – review and editing: **Aninda Arnab Pandit:** Methodology, **Joyanti Ray:** Writing – review and editing**: Md. Monirul Islam:** Supervision, Writing – review and editing:

## Supporting information

S1 Fig. Location of the experiment

S2 Fig. Manifested weather situation during crop growing period

S1 Table. Information about the selected 30 SSR markers used in this study.

S2 Table. Analysis of variance (mean square) for 17 characteristics of 100 mutants (M3) and their parent

S3 Table. Z score

S4 Table. Procrustes coordinates

S5 Table. Core Germplasm Network Analysis

## Reference

Al Mamun MA, Islam MM, Adhikary SK, Haque MM, Hasan MJ, Seraj ZI, et al. Genotype selection from azide-induced rice mutants using multitrait genotype-ideotype distance index (MGIDI): Unveiling promising variants for yield improvement. Adv Agric. 2024; 2024:5719580. doi: 10.1155/2024/5719580

Al-Daej MI, Rezk AA, El-Malky MM, Shalaby TA, Ismail M. Comparative genetic diversity assessment and marker–trait association using two DNA marker systems in rice (*Oryza sativa* L.). Agronomy. 2023;13(2):329. doi: 10.3390/agronomy13020329

Allier A, Teyssèdre S, Lehermeier C, Moreau L, Charcosset A. Optimized breeding strategies to harness genetic resources with different performance levels. BMC Genomics. 2020;21(1):349. doi: 10.1186/s12864-020-6756-0

Bandumula N. Rice production in Asia: Key to global food security. Proc Natl Acad Sci India Sect B Biol Sci. 2018;88(4):1323–1328. doi: 10.1007/s40011-017-0867-7

Bonkoungou TO, Adejumobi II, Adetimirin VO, Badu-Apraku B, Agre PA, Nanema KR, et al. Joint analysis of phenotypic and molecular data for genetic diversity assessment in extra-early orange maize (Zea mays L.). BMC Genomics. 2025; 26:784. doi: 10.1186/s12864-025-11964-5

Burton GW, DeVane EH. Estimating Heritability in Tall Fescue (Festuca Arundinacea) from Replicated Clonal Material. Agron J. 1953;45(10):478–481. doi: 10.2134/agronj1953.00021962004500100005x

Charlesworth B, Jensen J. Effects of selection at linked sites on patterns of genetic variability. Annu Rev Ecol Evol Syst. 2021; 52:177–197. doi: 10.1146/annurev-ecolsys-010621-044528

Chen H, Sawasdee A, Lin Y, Chiang M, Chang H, Li W, et al. Reverse mutations in pigmentation induced by sodium azide in the IR64 rice variety. Curr Issues Mol Biol. 2024;46(12):13328–13346. doi: 10.3390/cimb46120795

Chen J, Zhou H, Xie W, Xia D, Gao G, Zhang Q, et al. Genome-wide association analyses reveal the genetic basis of combining ability in rice. Plant Biotechnol J. 2019;17(11):2211–2222. doi: 10.1111/pbi.13134

Chen X, Temnykh S, Xu Y, Cho YG, McCouch SR. Development of a microsatellite framework map providing genome-wide coverage in rice (*Oryza sativa* L.). Theor Appl Genet. 1997;95(4):553–567. doi: 10.1007/s001220050596

Comstock RE, Robinson HF. Estimation of average dominance of genes. In: Gowen JW, editor. Heterosis. Ames: Iowa State College Press; 1952. pp. 494–516.

Das A, Sahoo DR, Barik D, Subudhi E. Identification of duplicates in ginger germplasm collection from Odisha using morphological and molecular characterization. Proc Natl Acad Sci India Sect B Biol Sci. 2020; 90:1149–1156. doi: 10.1007/s40011-020-01178-y

Debnath P, Chakma K, Bhuiyan MSU, Thapa R, Pan R, Akhter D. A novel multi-trait genotype-ideotype distance index (MGIDI) for genotype selection in plant breeding: Application, prospects and limitations. Crop Design. 2024;3(4):100074. doi: 10.1016/j.cropd.2024.100074

Deshmukh R, Sonah H, Singh VP, Nguyen HT. Two novel QTLs for the harvest index that contribute to high-yield production in rice (*Oryza sativa* L.). Rice. 2021; 14:14. doi: 10.1186/s12284-021-00456-1

Evanno G, Regnaut S, Goudet J. Detecting the number of clusters of individuals using the software STRUCTURE: a simulation study. Mol Ecol. 2005;14(8):2611–2620. doi: 10.1111/j.1365-294X.2005.02553.x

Gill M, DeSalvatore D, Adak A, Thompson A, Li M, Ratnaparkhe MB, et al. Machine learning models outperform deep learning models, provide interpretation and facilitate feature selection for soybean trait prediction. BMC Plant Biol. 2022; 22:180. doi: 10.1186/s12870-022-03559-z

Gruszka D, Szarejko I, Maluszynski M. Sodium azide as a mutagen. In: Shu QY, Forster BP, Nakagawa H, editors. Plant Mutation Breeding and Biotechnology. Vienna: FAO/IAEA; 2012. pp. 159–166. doi: 10.1079/9781780640853.0159

Gu Y, Li W, Jiang H, Wang Y, Gao H, Liu M, et al. Simultaneously improving grain yield and water and nutrient use efficiencies by enhancing the harvest index in rice. Plant Commun. 2023;4(4):100505. doi: 10.1016/j.xplc.2023.100505

Hasib KM, Ganguli PK, Kole PC. Evaluation of the performance of advanced generation lines of mutant x Basmati crosses of scented rice. J Int Acad. 2004;8(1):7–10.

Hill CB, Li Y, Brim-DeForest WB, Reeder LA, Al-Khatib K, Linquist BA, et al. Predictors of high rice yields in a high-yielding environment: Lessons from a yield contest. Field Crops Res. 2024; 319:109474. doi: 10.1016/j.fcr.2024.109474

Hoque A, Begum SN, Hassan L. Genetic diversity assessment of Rice (*Oryza sativa* L.) germplasm using SSR markers. Res Agric Livest Fish. 2015;1(1):37–46. doi: 10.3329/ralf.v1i1.22354

Hossain M, Jaim WMH, Alam MS. Impact of BRRI developed technologies on production, productivity and farm income. In: Saleque MA, editor. Annual Research Review 2012-13. Gazipur: Bangladesh Rice Research Institute; 2013.

Johnson HW, Robinson HF, Comstock RE. Estimates of genetic and environmental variability in soybeans and their implications in selection. Agron J. 1955;47(10):477–483. doi: 10.2134/agronj1955.00021962004700070009x

Karim MR, Ishikawa M, Ikeda M, Islam MT. Climate change model predicts 33% rice yield decrease in 2100 in Bangladesh. Agron Sustain Dev. 2012;32(4):821–830. doi: 10.1007/s13593-012-0096-7

Khush GS. Breaking the yield frontier of rice. GeoJournal. 1995;35(3):329–332. doi: 10.1007/BF00989140

Knapp SJ, Cole GS, Pincot DDA, Dilla-Ermita CJ, Bjornson M, Famula RA, et al. Transgressive segregation, hopeful monsters, and phenotypic selection drove rapid genetic gains and breakthroughs in predictive breeding for quantitative resistance to Macrophomina in strawberry. Hortic Res. 2024;11(2): uhad289. doi: 10.1093/hr/uhad289

Kumar A, Sharma D, Tiwari A, Jaiswal J, Singh N, Sood S. Genotyping-by-Sequencing analysis for determining population structure of finger millet germplasm of diverse origins. Plant Genome. 2016;9(2): plantgenome2015.07.0058. doi: 10.3835/plantgenome2015.07.0058

Linquist B, Al-Khatib K, Brim-DeForest W, Espe MB, Espino L, Leinfelder-Miles M, et al. Predictors of high rice yields in a high-yielding environment: Lessons from a yield contest. Field Crops Res. 2024; 322:109693. doi: 10.1016/j.fcr.2024.109693

Liu K, Muse SV. PowerMarker: an integrated analysis environment for genetic marker analysis. Bioinformatics. 2005;21(9):2128–2129. doi: 10.1093/bioinformatics/bti282

Lo K, Chen Y, Chiang M, Chen M, Panibe JP, Chiu C, et al. Two genomic regions of a sodium azide induced rice mutant confer broad-spectrum and durable resistance to blast disease. Rice. 2022;15(1):2. doi: 10.1186/s12284-021-00547-z

Mackay IJ, Cockram J, Howell P, Powell W. Understanding the classics: the unifying concepts of transgressive segregation, inbreeding depression and heterosis and their central relevance for crop breeding. Plant Biotechnol J. 2021;19(1):26–34. doi: 10.1111/pbi.13481

MacNish TR, Danilevicz MF, Bayer PE, Bestry MS, Edwards D. Application of machine learning and genomics for orphan crop improvement. Nat Commun. 2025;16(1):982. doi: 10.1038/s41467-025-56330-x

Marsh R, Delabre I, Dyke J, Antal M, Doherty B. Agri-food system resilience: A multi-capital conceptual framework. J Agric Food Syst Community Dev. 2021;10(2):1–21. doi: 10.5304/jafscd.2021.102.013

Mbebi AJ, Mercado F, Hobby D, Tong H, Nikoloski Z. Advances in multi-trait genomic prediction approaches: classification, comparative analysis, and perspectives. Brief Bioinform. 2025;26(3): bbaf211. doi: 10.1093/bib/bbaf211

McCouch SR, Teytelman L, Xu Y, Lobos KB, Clare K, Walton M, et al. Development and mapping of 2240 new SSR markers for rice (*Oryza sativa* L.). DNA Res. 2002;9(6):199–207. doi: 10.1093/dnares/9.6.199

Nachimuthu VV, Muthurajan R, Duraialaguraja S, Sivakami R, Pandian BA, Ponniah G, et al. Analysis of population structure and genetic diversity in rice germplasm using SSR markers: An initiative towards association mapping of agronomic traits in *Oryza sativa*. Rice. 2015;8(1):5. doi: 10.1186/s12284-015-0062-5

Nei M. Genetic distance between populations. Am Nat. 1972;106(949):283–292. doi: 10.1086/282771

Oladosu Y, Rafii MY, Abdullah N, Hussin G, Ramli A, Rahim HA, et al. Principle and application of plant mutagenesis in crop improvement: A review. Biotechnol Biotechnol Equip. 2016;30(1):1–16. doi: 10.1080/13102818.2015.1087333

Olivoto T, Nardino M. MGIDI: toward an effective multivariate selection in biological experiments. Bioinformatics. 2021;37(10):1383–1389. doi: 10.1093/bioinformatics/btaa981

Pallavi M, Maruthi Prasad BP, Shanthi P, Reddy VLN, Kumar AN. Multi-trait genotype-ideotype distance index (MGIDI) for early seedling vigour and yield related traits to identify elite lines in rice (*Oryza sativa* L.). Electron J Plant Breed. 2024;15(1):120–131. doi: 10.37992/2024.1501.020

Peakall R, Smouse PE. GenAlEx 6.5: genetic analysis in Excel. Population genetic software for teaching and research—an update. Bioinformatics. 2012;28(19):2537–2539. doi: 10.1093/bioinformatics/bts460

Pradhan SK, Barik SR, Sahoo A, Mohapatra S, Nayak DK, Mahender A, et al. Population structure, genetic diversity and molecular marker-trait association analysis for high temperature stress tolerance in rice. PLoS One. 2016;11(8): e0160027. doi: 10.1371/journal.pone.0160027

Pritchard JK, Stephens M, Donnelly P. Inference of population structure using multilocus genotype data. Genetics. 2000;155(2):945–959. doi: 10.1093/genetics/155.2.945

Reddy GE, Suresh BG, Sravan T, Reddy PA. Interrelationship and cause-effect analysis of rice genotypes in North East plain zone. Bioscan. 2013;8(4):1141–1144.

Ronald PC. Lab to farm: Applying research on plant genetics and genomics to crop improvement. PLoS Biol. 2014;12(6):e1001878. doi: 10.1371/journal.pbio.1001878

Salem KFM, Safhi FA, Alwutayd KM, Abozahra MS, Almohisen IAA, Alsharari SF, et al. Analysis of genetic diversity, population structure and phylogenetic relationships of rice (*Oryza sativa* L.) cultivars using simple sequence repeat (SSR) markers. Genet Resour Crop Evol. 2024;71(5):2213–2227. doi: 10.1007/s10722-023-01789-0

J. Sambrook, E.F. Fritsch, T. Maniatis. Molecular Cloning: A Laboratory Manual Cold Spring Harbor Laboratory, Cold Spring Harbor, NY (1989)

Seck PA, Diagne A, Mohanty S, Wopereis MCS. Crops that feed the world 7: Rice. Food Secur. 2012;4(1):7–24. doi: 10.1007/s12571-012-0168-1

Serrat X, Esteban R, Guibourt N, Moysset L, Nogués S, Lalanne E. EMS mutagenesis in mature seed-derived rice calli as a new method for rapidly obtaining TILLING mutant populations. Plant Methods. 2014; 10:5. doi: 10.1186/1746-4811-10-5

Shen J, Li Z, Chen J, Song Z, Zhou Z, Shi Y. SHEsisPlus, a toolset for genetic studies on polyploid species. Sci Rep. 2016; 6:24095. doi: 10.1038/srep24095

Sikora P, Chawade A, Larsson M, Olsson J, Olsson O. Mutagenesis as a tool in plant genetics, functional genomics, and breeding. Int J Plant Genomics. 2011; 2011:314829. doi: 10.1155/2011/314829

Singh A, Kumar D, Gemmati D, Ellur R, Singh A, Tisato V, et al. Investigating genetic diversity and population structure in rice breeding from association mapping of 116 accessions using 64 polymorphic SSR markers. Crops. 2024;4(2):180–194. doi: 10.3390/crops4020014

Singh A, Ganapathysubramanian B, Singh AK, Sarkar S. Machine learning for high-throughput stress phenotyping in plants. Trends Plant Sci. 2016;21(2):110–124. doi: 10.1016/j.tplants.2015.10.015

Singh N, Choudhury DR, Singh AK, Kumar S, Srinivasan K, Tyagi RK, et al. Comparison of SSR and SNP markers in estimation of genetic diversity and population structure of Indian rice varieties. PLoS One. 2013;8(12): e84136. doi: 10.1371/journal.pone.0084136

Tai TH, Chun A, Henry IM, Ngo KJ, Burkart-Waco D. Effectiveness of sodium azide alone compared to sodium azide in combination with methyl nitrosurea for rice mutagenesis. Plant Breed Biotechnol. 2016;4(4):453–461. doi: 10.9787/PBB.2016.4.4.453

Verma H, Borah JL, Sarma RN. Variability assessment for root and drought tolerance traits and genetic diversity analysis of rice germplasm using SSR markers. Sci Rep. 2019;9(1):16513. doi: 10.1038/s41598-019-52884-1

Viana VE, Pegoraro C, Busanello C, de Oliveira AC. Mutagenesis in rice: The basis for breeding a new super plant. Front Plant Sci. 2019; 10:1326. doi: 10.3389/fpls.2019.01326

Vinoth PK, Manonmani S, Raveendran M, Sritharan N, Sudha M, Puja M, et al. Comparison of traditional selection indices and the multi-trait genotype-ideotype distance index (MGIDI) for improving rice (*Oryza sativa* L.) cultivars. Plant Sci Today. 2025;12(sp1). Available from: https://horizonepublishing.com/index.php/PST/article/view/5475

Wang C, Yu X, Wang J, Zhao Z, Wan J. Genetic and molecular mechanisms of reproductive isolation in the utilization of heterosis for breeding hybrid rice. J Genet Genomics. 2024;51(6):583–593. doi: 10.1016/j.jgg.2024.01.007

Wang N, Long T, Yao W, Xiong L, Zhang Q, Wu C. Mutant resources for the functional analysis of the rice genome. Mol Plant. 2013; 6:596–604. doi: 10.1093/mp/sss142

Washburn JD, Cimen E, Ramstein G, Reeves T, O’Briant P, McLean G, et al. Predicting phenotypes from genetic, environment, management, and historical data using CNNs. Theor Appl Genet. 2021;134(12):3997–4011. doi: 10.1007/s00122-021-03943-7

Werner CR, Gaynor RC, Gorjanc G, Hickey JM, Kox T, Abbadi A, et al. How population structure impacts genomic selection accuracy in cross-validation: Implications for practical breeding. Front Plant Sci. 2020; 11:592977. doi: 10.3389/fpls.2020.592977

Williams, J.G.K., Kubelik, A.R., Livak, K.J., Rafalski, J.A. and Tingey, S.V. (1990). DNA Amplified by Arbitrary Primers are Useful Genetic Markers, Nucleic Acids Research, 18(22), pp. 6531–6535, Available at: 10.1093/nar/18.22.6531.

Yamamoto T, Suzuki T, Suzuki K. Assessment of agro-morphological, physiological and yield traits diversity among tropical rice. PeerJ. 2021;9: e11752. doi: 10.7717/peerj.11752

Yang J, Zhang J. Grain-filling problem in ‘super’ rice. J Exp Bot. 2010;61(1):1–5. doi: 10.1093/jxb/erp348

Yang J, Zhang J. Crop management techniques to enhance harvest index in rice. J Exp Bot. 2010;61(12):3177–3189. doi: 10.1093/jxb/erq112

Yang J, Zhang J. Simultaneously improving grain yield and water and nutrient use efficiencies by enhancing the harvest index in rice. Crop Environ. 2023;2(3):157–164. doi: 10.1016/j.crope.2023.07.001

Zhang H, Zhu YJ, Zhu AD, Fan YY, Huang TX, Zhang JF, et al. Improvements in grain yield and nutrient utilization efficiency of japonica inbred rice released since the 1980s in eastern China. Crop J. 2022;10(1):225–233. doi: 10.1016/j.cj.2021.04.009

